# Transgenerational transcriptional heterogeneity from cytoplasmic chromatin

**DOI:** 10.1101/2022.01.12.475869

**Authors:** Stamatis Papathanasiou, Nikos A. Mynhier, Shiwei Liu, Etai Jacob, Ema Stokasimov, Bas van Steensel, Cheng-Zhong Zhang, David Pellman

**Affiliations:** Department of Cell Biology, Blavatnik Institute, Harvard Medical School, Boston, MA, USA; Department of Pediatric Oncology, Dana-Farber Cancer Institute, Boston, MA, USA; Department of Chemistry and Chemical Biology, Harvard University, Cambridge, MA, USA; Single-Cell Sequencing Program, Dana-Farber Cancer Institute, Boston, MA, USA; Department of Data Sciences, Dana-Farber Cancer Institute, Boston, MA, USA; AstraZeneca, 35 Gatehouse Dr, Waltham, MA 02451; Division of Gene Regulation and Oncode Institute, the Netherlands Cancer Institute, Amsterdam, the Netherlands; Department of Biomedical Informatics, Blavatnik Institute, Harvard Medical School, Boston, MA, USA; Howard Hughes Medical Institute, Chevy Chase, MD, USA

## Abstract

Transcriptional heterogeneity from plasticity of the epigenetic state of chromatin is thought to contribute to tumor evolution, metastasis, and drug resistance ^1–3^. However, the mechanisms leading to nongenetic cell-to-cell variation in gene expression remain poorly understood. Here we demonstrate that heritable transcriptional changes can result from the formation of micronuclei, aberrations of the nucleus that are common in cancer^4,5^. Micronuclei have fragile nuclear envelopes (NE) that are prone to spontaneous rupture, which exposes chromosomes to the cytoplasm and disrupts many nuclear activities ^6,7^. Using a combination of long-term live-cell imaging and same-cell, single-cell RNA sequencing (Look-Seq2), we identified significant reduction of gene expression in micronuclei, both before and after NE rupture. Furthermore, chromosomes in micronuclei fail to normally recover histone 3 lysine 27 acetylation, a critical step for the reestablishment of normal transcription after mitosis ^8–10^. These transcription and chromatin defects can persist into the next generation in a subset of cells, even after these chromosomes are incorporated into normal daughter nuclei. Moreover, persistent transcriptional repression is strongly associated with, and may be explained by, surprisingly long-lived DNA damage to these reincorporated chromosomes. Therefore, heritable alterations in transcription can originate from aberrations of nuclear architecture.

## Main

Nuclear atypia, encompassing aberrations of nuclear size and morphology, is a hallmark feature of many tumors that is commonly used to assign tumor grade and predict patient prognosis^4,11^. Recently, our group and others determined that structural abnormalities of the nucleus–micronuclei or chromosome bridges–can lead to a variety of simple and complex chromosomal rearrangements, including an extensive form of chromosome rearrangement called chromothripsis^12–15^. Chromothripsis enables rapid genome evolution of many cancers^16,17^. Although the role of nuclear abnormalities in self-amplifying genetic instability is now appreciated, other consequences of nuclear atypia have been little studied. For example, although micronuclei have been reported to have transcription defects and altered chromatin marks^6,18,19^, the functional consequences of these alterations remain poorly understood.

### Look-Seq2, live-cell imaging and same-cell RNA sequenching

A direct assessment of the transcriptional consequences of micronucleation requires single-cell analysis. We extended our previous method of live-imaging and single-cell sequencing^13,20^ to perform single-cell RNA sequencing on micronucleated cells and their family members. We induced chromosome missegregation and generated micronucleated cells as previously described^13^. We then assessed chromosome dynamics and the maintenance or loss of micronuclear NE integrity by live-cell imaging (Supplementary Video 1). Cells were then isolated for single-cell RNA-Seq (scRNA) library construction by SMARTSeq2^21^(Extended Fig. 1a). Initially, cell isolation was performed similarly to our approach for single-cell whole genome sequencing^13,20^. However, for most of the experiments in this study, we used an improved contact-free laser capture microdissection method (Extended Fig. 1b and Methods) that is not only optimized for isolating cells with minimal perturbation, but also facilitated the isolation of micronucleated cell’s daughters, sister, or nieces (in experiments where the sister divided). Hereafter, we refer to this method, with either cell capture technique, as “Look-Seq2”.

The computational analysis Look-Seq2 data needs to accomplish two goals. First, we need to identify the chromosome in the micronucleus (“generation 1”, Extended Fig. 1a) or the chromosome that was in the micronucleus in the first generation and then reincorporated into a normal daughter nucleus in the second generation (“generation 2”). Second, we need to quantitatively assess the transcription output from this chromosome (transcription yield) relative to its homologue and the other chromosomes in the main nucleus (Extended Fig. 1c). To achieve these goals, we need to determine both the segregation pattern of the micronuclear chromosome in both generations and the parental haplotype of the micronuclear chromatid to measure haplotype-specific expression.

We designed an analytical workflow with three major components (Methods, Extended Data Fig. 2). First, we generated scRNA data for a large number of isogenic RPE-1 cells as a reference to estimate both average gene expression and the range of spontaneous transcriptional variation unrelated to micronucleation in disomic chromosomes (Extended Data Fig. 1c,d and Supplementary Table 1, Methods). This enabled us to measure the transcription yield from all chromosomes in the micronucleated cell’s family and identify chromosomes with non-disomic transcription output. Second, based on the patterns of total and allelic expression in each cell family, we determined the integer copy number ratio of transcribed chromosomes (Methods, Extended Data Fig. 2 and Supplementary Table 2). Because only two possible segregation patterns can generate the micronucleated cell, and because transcription in the normal nucleus of the sister cell (or its progeny) directly reflects the underlying DNA copy number, the segregation pattern and the micronuclear haplotype can be identified independent of assessing the transcription output of the chromosome from the micronucleus. As an example, trisomy in a micronucleated cell (2C+1) can be identified because this cell’s sister (or nieces) will have monosomic transcription for that chromosome due to the segregation pattern generating the trisomy (Fig. 1a, 1:3 segregation), regardless of whether the micronuclear chromosome underwent complete silencing (resulting in a normal diploid transcriptome) or was normally transcribed (resulting in a trisomic expression). Finally, we used the parental haplotype information of RPE-1 cells^13,20,22^ to verify the conclusions about the segregation patterns and transcription yield from the micronuclear chromosome (Methods and Supplementary Table 2).

**Figure 1.**
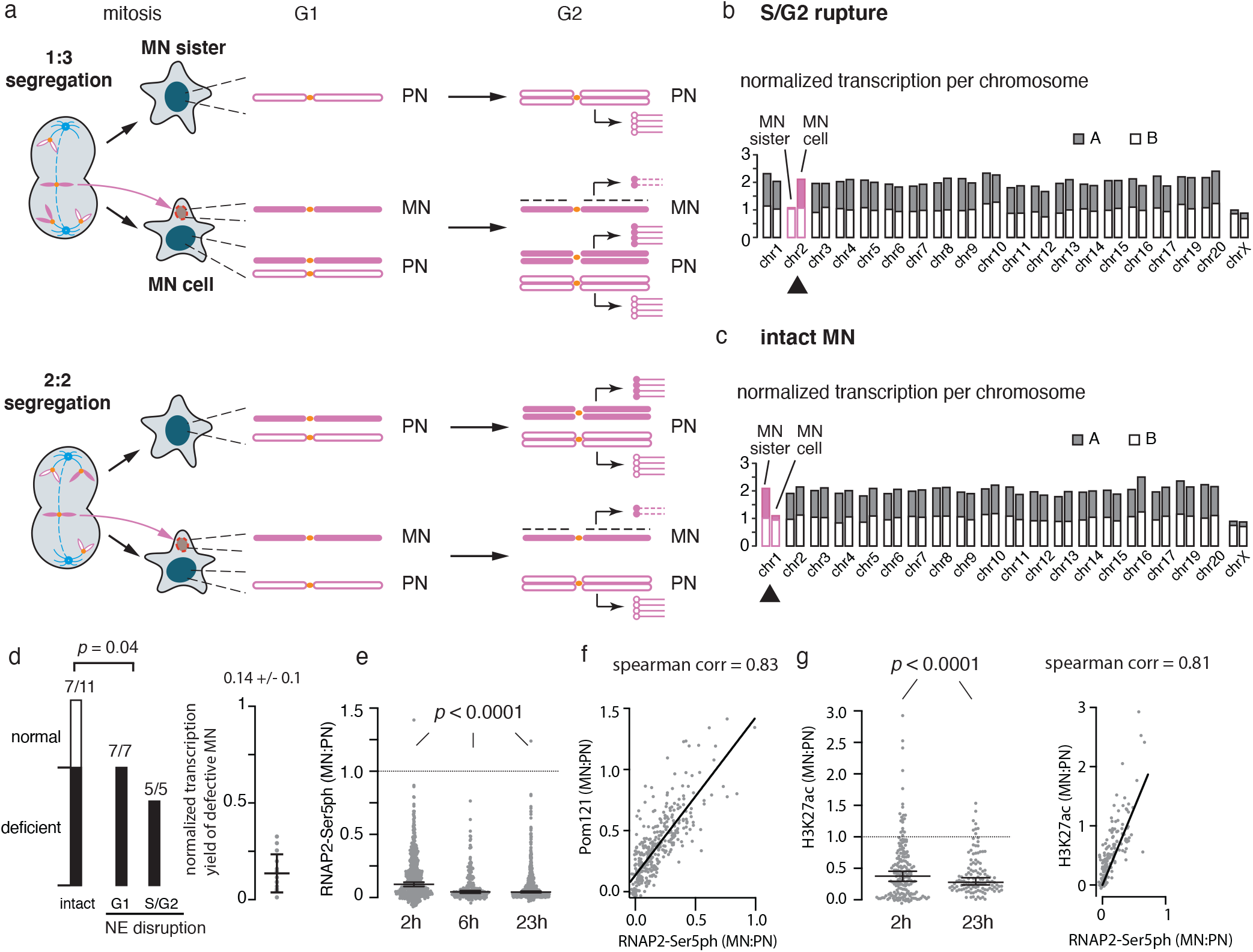
Transcription defects in newly-generated micronuclei. (a) Two segregation patterns of lagging chromosomes in the micronucleated cell (“MN cell”) and its sister (“MN sister”). The lagging chromosome (filled) and its homolog (open) are colored in magenta, other chromosomes are colored gray. Top, under 1:3 (mis)segregation, the G1 sister is monosomic and the G1 MN cell is trisomic; Bottom, under 2:2 segregation, both the MN cell and the sister are disomic. In G2 (when cells were isolated), chromosomes in the primary nucleus (PN) are replicated but the chromosome in the micronucleus (MN) is poorly replicated. Transcripts (lollipops) from each chromatid (open circles for the normally segregated homolog, filled circles for the MN homolog, and dashed lines for transcripts from the MN chromatid) are normalized against parental RPE-1 cells to calculate the average transcription yield. (b) Chromosome-wide transcription silencing of a MN generated by a 1:3 segregation that underwent NE rupture in S/G2 (a, top). Shown is normalized total transcription from both homologs and the allelic ratio of all chromosomes except Chrs.18, 21, and 22. Gray and white represent contributions from two parental homologs (A and B) determined from the parental haplotype. The micronuclear chromosome (filled) and the other homolog (open) are shown in magenta as in (a). (c) Chromosome-wide silencing of an intact MN generated by a 2:2 segregation (a, bottom). In contrast to (b), there was disomic expression in the MN cell. (d) Summary of the frequency of transcription defects in 23 pairs of MN cell/MN sister (Generation 1). The numbers of cells with or without transcriptional deficiency are shown for cells with intact MN, cells that underwent MN NE rupture in G1 (before 8 hours post mitotic shake-off), and cells that underwent MN NE rupture in S/G2 (after 8 hours post-mitotic shake-off). The average transcription yield of defective MNs relative to normal transcription (estimated from 2:2 segregation examples) is 0.14 +/- 0.1 (14 total MN chromosomes from 13 families). The p-value is from one-sided Fisher’s exact test. (e) Quantitative analysis of the RNAP2-Ser5ph signal shows transcription defects in all MN, irrespective of the timepoint during interphase at which transcription was assessed. Values are the MN:PN ratio of fluorescence intensity (FI) units (n = 644, 212 and 695, left to right, from at least three experiments). Dotted line, the expectation for normal MN transcription (MN:PN FI ratio 1). Shown is the median ratio with a 95% confidence interval (CI). Two-tailed Mann–Whitney test (Extended Data Fig. 1b). (f) MN transcription is correlated with the density of NPCs (POM121) at 2 h post mitotic shake-off (n = 225, three experiments). Two-tailed Spearman’s correlation. (g) MN chromosomes exhibit decreased H3K27ac. Left, MN:PN ratios for H3K27ac, as in (e). Right, Correlation between H3K27ac and RNAP2-Ser5ph MN:PN ratios (n = 187, two experiments).

### Transcriptome defects from newly generated micronuclei

To evaluate the specificity of Look-Seq2, we initially focused on cells with NE ruptured micronuclei that were expected to display near-complete transcriptional silencing^6^. We first analyzed acentric micronuclear chromosomes generated by CRISPR-Cas9 cleavage of the q arm of Chr 5 during generation 1 (Extended Data Fig. 3a)^23^. In these samples, because the identity of the micronuclear chromosome arm was predetermined by Cas9-cleavage, transcriptional silencing should be unambiguously identifiable from the analysis of sister pairs. Indeed, we identified the transcriptional patterns expected from silencing of the Cas9-generated acentric fragment, along with the expected underlying segregation patterns inferred from the sister cell transcriptomes (Leibowitz et al.^23^ and Extended Data Fig. 3a,b).

Next, we generated micronucleated cells by random whole chromosome missegregation and analyzed the transcriptomes of 12 of these cells that underwent micronuclear NE disruption. Our analysis of these cells and their sisters identified the micronuclear chromosomes that underwent near-complete transcriptional silencing aftern NE disruption and distinguished 1:3 or 2:2 segregations patterns (Fig.1a, Extended Data Fig. 2a, and Supplementary Table 2). In one example shown in Fig.1b and Extended Data Fig. 3c, only a single Chr. 2 homolog displayed the expected pattern of allelic imbalance and was inferred to be the chromatid in the micronucleus; all other chromosomes displayed normal bi-allelic disomic transcription except Chr.X that displayed shared monoallelic expression reflecting X-chromosome inactivation in female RPE-1 cells. Together, these results demonstrate that Look-Seq2 can identify chromosome arm-level gene expression silencing through haplotype-resolved transcriptome analysis.

We next used Look-Seq2 to determine if chromosomes in intact micronuclei exhibit transcriptional defects (Methods). Qualitative analysis suggested that transcription might occur at normal levels in intact micronuclei^6^; however, this is at odds with other work indicating that many nuclear functions are compromised in micronuclei, at least in part due to their defective nucleocytoplasmic transport^24^. Consistent with the later results, we found that most intact micronuclei also exhibited strong transcriptional defects (Fig. 1c and Extended Data Fig. 3d), although a small fraction of intact micronuclei appeared to be normally transcribed (Fig. 1d, p = 0.04, one-sided Fisher’s exact test). In support of this conclusion, fluorescence intensity (FI) measurements indicated a significant decrease in a marker of active, phosphorylated RNA Polymerase II (RNAP2-Ser5ph) in both intact and ruptured micronuclei relative to their primary nuclei (Extended Data Fig. 4a-e). Notably, this transcription defect was evident from the beginning of interphase (Fig. 1e and Extended Data Fig. 4b,c) and the degree of RNAP2-Ser5 loss is positively correlated with the extent of the defect in nuclear pore complex assembly (Fig. 1f), which our prior studies demonstrated is itself correlated with defects in nuclear import^7,24^.

We finally assessed whether the transcriptional defects of micronuclear chromosomes were correlated with alterations in epigenetic chromatin marks. We observed a modest increase in repressive heterochromatin marks histone 3 lysine 9 dimethylation (H3K9me2) or histone 3 lysine 27 trimethylation (H3K27me3) that accumulated on a subset of micronuclei with NE disruption late during interphase (Extended Data Fig. 4f, g), consistent with a previous report^18^. More strikingly, we found that micronuclei exhibited a significant loss of the active chromatin marks histone 3 lysine 27 acetylation (H3K27ac) and histone 3 lysine 9 acetylation (H3K9ac)^18^ from the beginning of interphase that was positively correlated to the level of active RNA Pol II (Fig. 1g and Extended Data Fig. 4h). This highly penetrant loss of H3K27ac is notable because recent studies indicated that recovery of H3K27ac is essential for the normal reestablishment of transcription after mitosis^8–10^.

In summary, both single-cell RNA-Seq and cell biological analyses demonstrated that chromosomes in micronuclei acquire profound transcriptional defects that are not solely due to micronuclear NE rupture. The correlated acquisition of altered chromatin states raises the possibility that the transcription defects could be inherited.

### Micronuclear chromosomes acquire heritable transcription defects

After cell division, micronuclear chromosomes either remain in the cytoplasm of a daughter cell, separate from the main nucleus or, if positioned close enough to the main mass of chromosomes during mitosis, can be incorporated into newly formed daughter cell nuclei (“generation 2”, Fig. 2a and Extended Data Fig. 1a). To determine if transcription defects of micronuclear chromosomes persist even in a normal nuclear environment, we performed Look-Seq 2 analysis on daughters of micronucleated cells (MN daughters) after micronuclear chromosomes were incorporated into normal nuclei after cell division. The average time interval after reincorporation for these samples was 16.1 hours, which far exceeds the time required for normal chromosomes to recover transcription after mitosis (90 min)^8–10^. We observed heterogeneous transcriptional recovery of the micronuclear chromosome after reincorporation. The identity and segregation pattern of the micronuclear chromosome was determined as described above (Fig. 2a, Extended Data Fig. 2c and Extended Data Fig. 5; Methods, Supplementary Table 2). Whereas a majority of reincorporated chromosomes from micronuclei exhibited normal post-mitotic transcriptional recovery (Supplementary Table 2), a subset of reincorporated micronuclear chromosomes exhibited a significant reduction or near complete loss of transcription (11/38 samples, Fig. 2a-c, Extended Data Fig. 5, Extended Data Fig. 6 and Supplementary Table 2). The examples of near-complete silencing of the reincorporated micronuclear chromosome were important, because, unlike partial silencing, chromosome-wide silencing cannot be explained by segmental chromosome loss associated with chromothripsis (Extended Data Fig. 6c).

**Figure 2.**
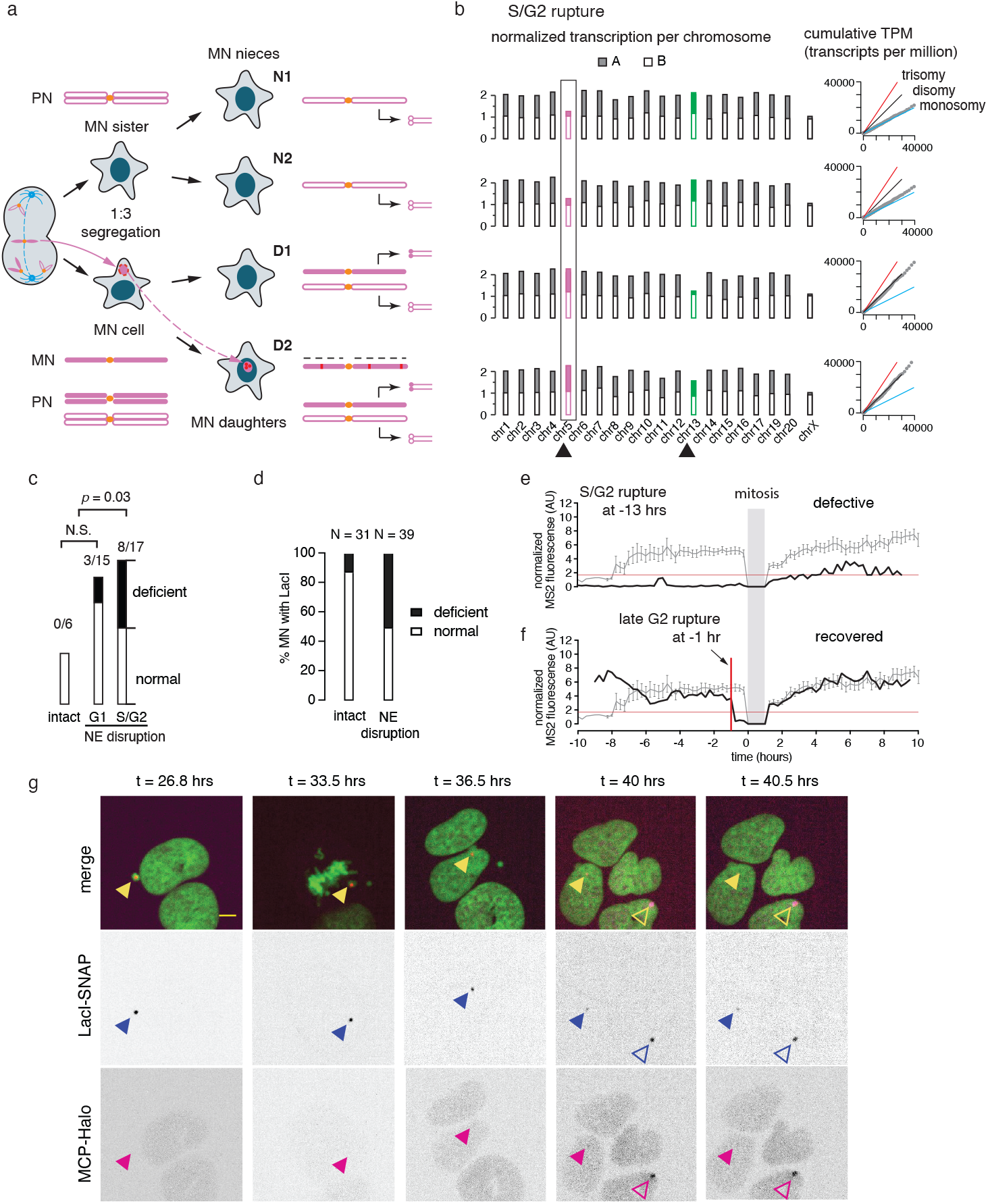
Variably penetrant transcription compromise after MN chromosome reincorporation into a normal nucleus. (a) The expected copy-number outcome and transcriptional yield of progeny cells after two generations following a 1:3 segregation in the first generation (Fig. 1a, top). For the 2:2 segregation, see Extended Data Fig. 5c. The monosomic MN sister cell gives rise to two monosomic “nieces” of the MN cell (MN nieces); division of the MN cell gives rise to two daughters (MN daughters). Shown is the G1 chromosome copy number for nieces and daughters. Because the MN chromosome is poorly replicated during generation 1, the MN daughter that reincorporates the MN chromosome should be trisomic, whereas the other MN daughter should be disomic. The pattern of monosomic transcription in both MN nieces and disomic transcription in at least one MN daughter cell establishes the missegregation pattern and identify the MN chromosome. (b) Persistent loss of transcription of a reincorporated MN chromosome (Chr. 5) after NE rupture in S/G2 and reincorporation to the daughter MN. Left, the normalized total and allelic transcription yield (filled and open bars) of both nieces and daughters. Two chromosomes (Chr.5 and Chr.13) display transcriptional imbalance consistent with 1:3 segregation in the first generation (Chr.5, magenta) and 2:2 segregation (Chr.13, green). Right, observed cumulative transcription yield (from low to high expression) on Chr.5 relative to expectations for monosomic (blue), disomic (black), and trisomic (red) expression. In monosomic MN cell nieces, monosomic transcription is evident from the reduced total transcription and the loss of MN haplotype transcripts. By contrast, both MN daughters display close to disomic total transcription and 1:1 allelic ratios, indicating that there is no measurable increase in transcription from the reincorporated copy of Chr.5 from the micronucleus (which MN daughter has the extra, silenced copy of Chr.5 cannot be determined). For Chr.13, the bottom cell (D2) displays partial transcription of the MN haplotype and was inferred to be the cell of Chr.13 reincorporation (see Extended Data Fig. 5). (c) Summary of the transcription of reincorporated MN in 38 families of MN daughters and nieces. Both p-values are from one-sided Fisher’s exact test. Deficient transcription recovery is separately assessed for 1:3 segregations and 2:2 segregations. For 1:3 segregation, deficient transcription recovery is defined as when the ratio of total transcription between MN daughters is significantly lower than 1.5:1 as expected from the DNA copy number ratio after reincorporation (3:2). For 2:2 segregation, deficient transcription recovery is defined as when the relative transcription yield of the MN haplotype in the daughter of MN reincorporation is significantly below 1. (d) Summary of experiments with the U2OS 2-6-3 transcription reporter to assess transcription from reincorporated MN chromosomes from generation 2 (n = 70 reincorporated chromosomes from 13 experiments). “Defect” in these experiments was no visible MS2 signal detected by the MCP-Halo tracer. Fisher’s exact test. (e) Example of generation 2 MN chromosome transcription defect shown by fluorescence intensity measurements of the MS2 signal over time. Grey line shows the average and SEM of controls where the reporter was in a normal nucleus before and after mitosis. Red horizontal line: minimum detectable value in the controls. Black line shows an example where the MN ruptured in S/G2 (not shown). MS2 expression is defective before and after mitosis. (f) Similar to (d), example of transcription recovery in normal levels after MN reincorporation. Red vertical line: time of MN rupture. (g) Images from a timelapse series showing defective transcription from the U2OS 2-6-3 reporter after MN rupture in S/G2 and reincorporation into a normal daughter nucleus. Shown are single focal plane confocal images at timepoints (hours) after release from a mitotic block. Green: GFP-H2B. Blue arrowheads: reporter chromosome detected by LacI focus; in generation 1 in micronucleus and in generation 2 is reincorporated into a daughter nucleus. Filled magenta arrowheads: MS2 expression of the locus. Open magenta arrowheads: neighboring cell that enters the field of view and serves as a MS2 detection control. The intensity plot for this cell is presented in (e). Scale bar 5 µm.

A closer inspection of the combined data for complete and partial transcriptional repression (Fig. 2c) revealed that NE rupture and its cell cycle timing influenced the extent of transcriptional recovery. If micronuclei in the first generation either formed without an intact NE, ruptured within the approximate time interval of G1, or remained intact throughout interphase, almost all micronuclear chromosomes recovered transcription in the second generation after incorporation into a normal daughter nucleus (18/21 samples, Fig. 2c, Extended Data Fig. 6a and Supplementary Table 2). By contrast, when micronuclei ruptured in the S/G2 interval of generation 1, transcriptional defects were detected in 8/17 samples (Fig. 2c, Extended Data Fig. 6b and Supplementary Table 2). The frequency of transcriptional deficiency of reincorporated micronuclei with S/G2 NE disruption (8/17) is significantly higher than the frequency of deficient reincorporated micronuclei with intact NE or early NE disruption (p = 0.04, one-sided Fisher’s exact test). In summary, Look-Seq2 experiments suggest that a subset of micronuclear chromosomes acquire heritable transcription defects that may be related to the cell cycle time during which NE rupture of the micronucleus occurs. We developed single-cell imaging approaches to verify and further study this phenomenon.

### Imaging nascent transcripts from micronuclear chromosomes

We adapted the U2OS 2-6-3 nascent transcription reporter system^25^ to independently assess the transcriptional activity of reincorporated micronuclear chromosomes. The 2-6-3 transcription reporter construct contains lac operator arrays, enabling visualization of the reporter locus, as well as an inducible mRNA containing MS2 aptamers, enabling the visualization of inducible nascent transcripts. We induced micronucleation in this cell line and cells with micronuclei containing the reporter were identified. After the division of these micronucleated cells, we analyzed generation 2 daughter cells where the chromosome with the reporter construct was reincorporated into a normal nucleus. Transcriptional activity of the reporter locus was assessed, either qualitatively, by the presence or absence of the MS2-marked transcript (Fig. 2d, Supplementary Video 2), or quantitatively, by measuring the fluorescence intensity of nascent transcripts from time-lapse confocal images using an automated analysis pipeline (Fig. 2e,f and Extended Data Fig. 7, see Methods). Consistent with the Look-Seq2 analysis, our imaging of nascent transcripts confirmed that a subset of reincorporated micronucleated chromosomes (24/70) exhibit persistent defects in transcription (Fig. 2d,e,g and Extended Data Fig. 7). Moreover, this analysis confirmed that 83% (20/24) of examples exhibiting a second-generation transcriptional defect underwent rupture of the micronucleus NE in the interphase of the prior cell cycle (Fig. 2d,e,g and Extended Data Fig. 7b, Supplementary Video 2).

### Heritable transcription defects originate from damaged chromosomes

NE rupture of micronuclei during the S/G2 phase of the cell cycle is associated with DNA damage more strongly than loss of NE integrity in G1^6,26^. Moreover, previous studies have shown that DNA damage response signaling triggers transcriptional silencing^27–29^. We therefore considered the possibility that the subset of micronuclear chromosomes exhibiting transcriptional suppression after reincorporation into a normal nucleus corresponds to the subset of micronuclear chromosomes that experience extensive DNA damage.

As a first test of this hypothesis, we used our previously described correlated live-cell same-cell fixed imaging protocol^20,24^ to follow micronuclear chromosomes through cell division, observe their reincorporation into a normal nucleus, and detect *γ*H2AX-marked DNA damage by immunofluorescence (hereafter “same-cell live-fixed” imaging, see Methods). From live cell imaging of GFP-H2B, we followed the division of 13 micronucleated cells that likely had reincorporation of the micronuclear chromosome because neither daughter had detectable micronuclei. In 8/13 of these cell divisions, we observed large *γ*H2AX-labeled subnuclear territories that were typically restricted to one of the two daughter nuclei (Extended Data Fig. 8a). We term these structures “MN-bodies”.

To determine if these *γ*H2AX-labeled MN-bodies are derived from reincorporated micronuclear chromosomes, we generated a fluorescent protein that specifically labeled damaged micronuclear chromosomes. We fused a DNA damage response protein which binds *γ*H2AX (MDC1, Mediator Of DNA Damage Checkpoint 1) with a tag that can be visualized with a fluorescent dye (SNAP-tag). During live-cell imaging, the SNAP-MDC1 fusion was not visible on micronuclear chromosomes during interphase, likely because it is sequestered in the main nucleus. However, upon mitotic NE breakdown, micronuclear chromosomes were brightly labeled, enabling them to be tracked through from mitosis into the next interphase (Fig. 3a and Supplementary Video 3). After division, a substantial fraction of these chromosomes (31/69) were incorporated into normal daughter nuclei to become nuclear MN-bodies (Fig. 3a and Extended Data Fig. 8b). Unexpectedly, the DNA damage detected in MN-bodies persisted for an extended time period (average of at least 20.8 hr, Extended Data Fig. 8c), longer than the normal time course of DNA double strand break repair^30^. 21/31of damaged MN-bodies were derived from micronuclei that ruptured during the prior interphase, suggesting that DNA damage was acquired at the time of micronuclear NE rupture and then carried over into MN-bodies formed in the second generation (Extended Data Fig. 8b). The fact that micronuclei can acquire DNA damage in mitosis^20^, as well as after interphase NE rupture, may explain why a minority of micronuclei that do not rupture in interphase can still generate MN-bodies with DNA damage.

**Figure 3.**
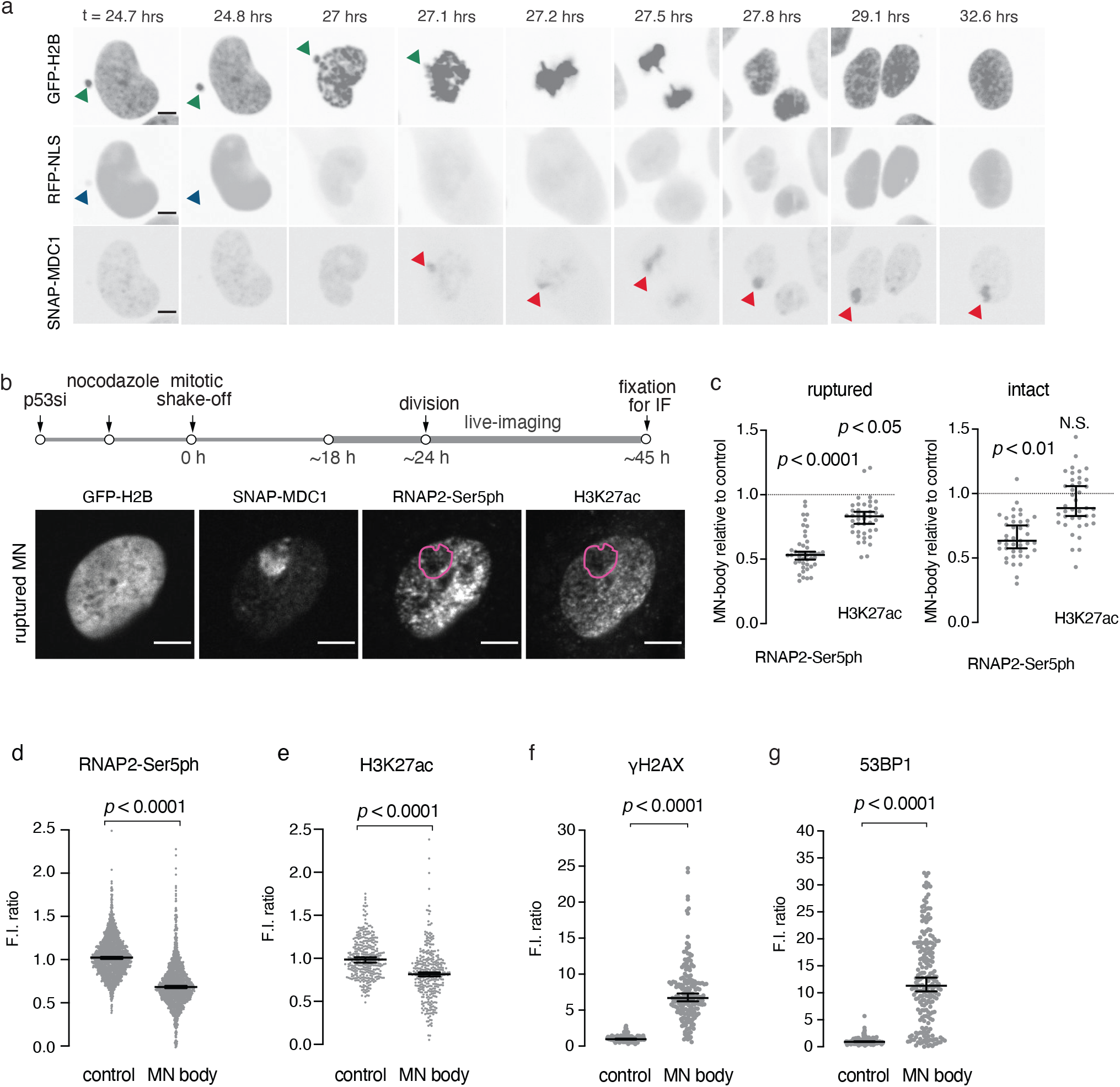
MN-bodies exhibit transcription defects and extensive DNA damage. (a) Tracking damaged MN chromosomes through and MN-body formation. Shown are images from a timelapse series. GFP-H2B: chromosomes; green arrowhead: MN chromosome; RFP-NLS: NE integrity; blue arrowheads: MN NE rupture; red arrowheads: SNAP-MDC1-marked MN DNA damage. Time: hours post release from the mitotic block for MN induction. Scale bars 5 µm. (b) Defective transcription and H3K27ac in MN-bodies. Top, scheme of the experiment (timepoints not to scale). Bottom, representative images of daughter cell with an MN-body from a ruptured MN. Magenta dashed line: MN-body with low RNAP2-Ser5ph and low H3K27ac. Scale bars 5 µm. (c) Aggregate relative MN body FI for RNAP2-Ser5ph and H3K27ac for experiments as in (b). Median with 95% CI; two-tailed Mann–Whitney test; comparing the FI units between MN and PN in the same cell. (d) Decrease of RNAP2-Ser5ph in MN-bodies verified by a large sample-size fixed imaging experiment. Cells are fixed at approximately 45 h post mitotic shake-off. MN-bodies were identified based on the SNAP-MDC1 signal. Relative FI measurements of MN-body RNAP2-Ser5ph (n = 1447, six experiments). Median with 95% CI; two-tailed Mann–Whitney test. (e) Decrease of H3K27ac in MN-bodies (n = 341, from two experiments). Analysis as in (d). (f) DNA damage in MN-bodies. FI measurements of *γ*H2AX intensity (94% of MN-bodies were positive; >3SD above background) for *γ*H2AX; n = 195, from two experiments). (g) 53BP1accumulation within MN-bodies (82% of MN-bodies were positive for 53BP1; n = 211, from two experiments).

We next used same-cell live-fixed imaging to determine if MN-bodies co-localized with markers of active transcription. Indeed, damaged MN-bodies exhibited reduced levels of both RNAP2-Ser5ph and H3K27ac (Fig 3b-c). These findings were confirmed by co-staining for these transcription markers with endogenous MDC1 (Fig. 3d-e and Extended Data Fig. 8d-f). MN-bodies not only accumulate *γ*H2AX and MDC1, but also accumulate the DNA damage response protein, 53BP1 (94% of MN bodies were positive for *γ*H2AX and 82% were positive for 53BP1, Fig. 3f,g and Extended Data Fig. 8d).

Although we and others have observed a modest accumulation of the repressive histone marks H3K9me2 and H3K27me3 in ruptured micronuclei^18^ (Extended Data Fig. 4f-g), our imaging show only a small, though significant, increase of these marks in MN-bodies (Extended Data Fig. 8g). Therefore, the loss of H3K27ac and RNAP2-Ser5ph appears to be the primary features associated with heritable transcriptional defects of micronuclear chromosomes.

The above data show that damaged chromosomes acquire transcriptional defects, however, they do not address whether it is primarily damaged chromosomes that acquire this defect, which would suggest that DNA damage and altered transcription could be mechanistically linked. Testing this association necessitated a live-cell imaging system that could track all micronuclear chromosomes, irrespective of whether they are damaged or not. To accomplish this, we generated a chromatin tagging system that we refer to as DamMN. DamMN is based on the ability of Dam (DNA-adenine methyltransferase) to methylate adenine residues in DNA resulting in N6-methyladenine (m6A)^31^. We generated an inducible Dam methyltransferase fused to three tandem copies of mCherry (“mega-Dam”, Fig. 4a and Extended Data Fig. 9a-c). Because we engineered the fusion to contain two tandem degrons, in synchronized cells, we could restrict its expression to the interphase when micronuclei formed by repressing its transcription and inducing fusion protein degradation just prior to mitotic entry. Because these manipulations result in the loss of mega-Dam prior to mitotic entry, this experimental design enables, in a large fraction of cells, specific labeling of the micronuclear chromosome without labeling the main chromosomes at the time of mitotic NE breakdown (Fig. 4a,b and Extended Data Fig. 9a-d).

**Figure 4.**
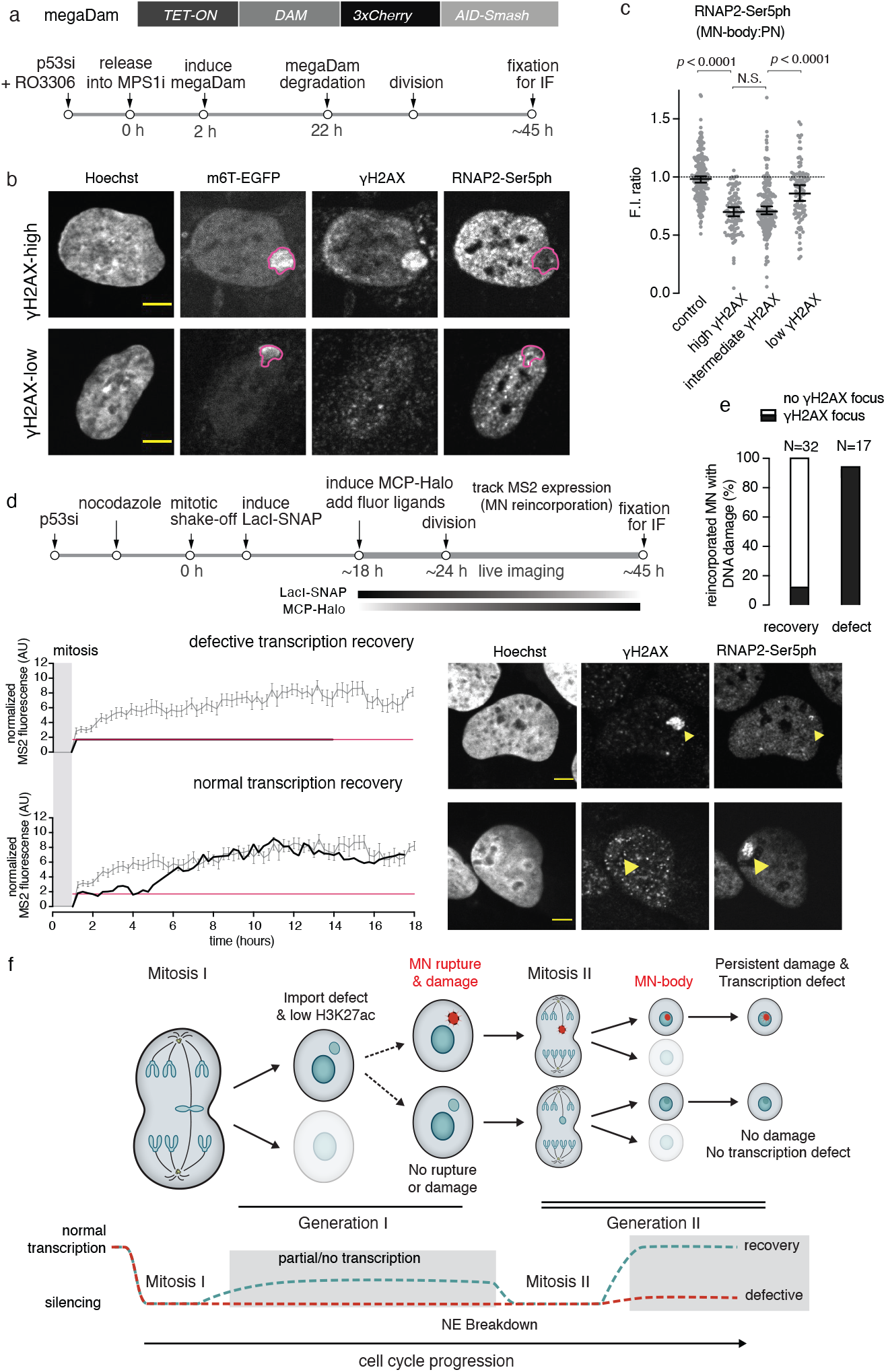
Damaged MN-bodies are more likely to have persistent transcription defects. (a)Transgenerational tracking of MN chromosome fate. Left top, cartoon of the gene expressing megaDam. Left bottom, Scheme of manipulations that restrict DamMN expression to the first interphase where the MN form. (b) Transcription state of high and low damage MN-bodies. Top, Representative image of a reincorporated MN marked by the m6A-Tracer with high (top quartile) *γ*H2AX-signal and low RNAP2-Ser5ph-labeling. Bottom, same but with low *γ*H2AX-signal (bottom quartile) and normal levels of RNAP2-Ser5ph-labeling. Scale bars 5 µm. Magenta dashed lines: MN-body area. (c) Quantification of aggregate data from (b) for the indicated samples (n = 220, 111, 220 and 112, left to right, four experiments). Median with 95% CI; Kruskal-Wallis with Dunn’s multiple comparisons test. (d) Correspondence between high damage level and low transcriptional activity for reincorporated MN chromosomes using the U2OS 2-6-3 nascent transcription reporter. Top: scheme of the experiment. During live imaging, cells with the reporter in a MN were identified by LacI-SNAP, followed through cell division to identify MN-body formation. Transcription activity of the reporter was tracked in live cells (MCP-Halo foci) and, after reincorporation, was scored for *γ*H2AX-marked DNA damage. Bottom left: MS2 FI signal used to detect reporter transcription activity, as in Fig. 2e. Bottom right: After live imaging, cells were fixed to detect *γ*H2AX, and RNAP2-Ser5ph (which correlated with the MS2 signal). Shown are representative images. Yellow arrowheads: MN body. Scale bars 5 µm. (e) Summary data for 29 cells with MN reincorporation as in (d). 29 cells with MN rupture during generation 1 and 20 cells with intact MN throughout interphase (from 11 experiments). The correlation between a transcription defect and DNA damage is significant (P<0.0001, Fisher’s exact test). (f) Top, scheme of cellular events leading to heritable transcription defects. Mitosis I: A cell with a lagging chromosome divides, generating the “MN cell” and its “sister” (greyed out). The MN NE can rupture (top, red) or remain intact (bottom). Mitosis II: The MN chromosome can be reincorporated into a normally functioning daughter nucleus (shown). MN rupture during S/G2 phase of the first cell division causes extensive DNA damage that persists into the next cell cycle, resulting in MN-bodies with varying degrees of transcriptional silencing (top, with red MN-body). Alternatively, undamaged or less damaged chromosomes will recover transcription after MN chromosome enters the normal daughter nucleoplasm (bottom). Bottom, two transcription outcomes for the chromosomes shown (top). Dotted aqua blue line: MN has an intact NE in generation 1 and recovers transcription in generation 2. Dotted red line: MN ruptures during generation 1, generating DNA damage that persists into generation 2 and fails to recover transcription after reincorporation.

We used the DamMN system to track micronuclear chromosomes through mitosis into the next interphase when a fraction of them formed MN-bodies. This analysis revealed that the MN-bodies with top quartile of *γ*H2AX labeling acquired a heritable defect in transcription, whereas the MN-bodies from the bottom quartile mostly recovered transcriptional activity (Fig. 4b,c and Extended Data Fig. 9e). We orthogonally validated this result with same-cell live-fixed imaging to detect *γ*H2AX and nascent transcription from the U2OS 2-6-3 system reporter. We found that 28/32 cells that recovered transcription lacked detectable DNA damage after generation 2 micronuclear chromosome reincorporation. By contrast, 16/17 of the chromosomes that exhibited persistent transcriptional suppression exhibited extensive DNA damage (Fig. 4d,e). Together, these results suggest a linkage between DNA damage and heritable transcriptome defects of micronuclear chromosomes.

## Discussion

Collectively, our data establish that micronuclei, a common feature of cancer nuclear atypia, can generate heritable defects in transcription. These findings should have relevance for tumor evolution^1–3^ as well as for contexts during normal development where micronucleation occurs, such as during epigenetically programed genome elimination^32^. We propose the following model for the acquisition of these heritable defects (Fig. 4f). When micronuclei form, even prior to NE rupture, they exhibit defects in post-mitotic transcriptional recovery along with variably reduced H3K27 acetylation. This initial transcriptional recovery defect likely stems from the abnormal composition of the nucleoplasm in micronuclei that results from their nuclear transport defects^7,24^. The lower levels of H3K27 acetylation persist after rupture of the micronucleus NE. However, multi-generational tracking showed that, in the next generation, these transcription defects can be reversible, unless they are accompanied by extensive DNA damage from micronuclear NE rupture during the interphase of the first cell cycle. Persistent DNA damage likely has a direct role in repressing transcription of micronuclear chromosomes because prior work has established that DNA damage or abnormal DNA replication generates transcriptional silencing and/or epigenetic plasticity^27,33^.

We note intriguing similarities between MN-bodies and previously described 53BP1 bodies^34–36^. 53BP1 bodies form during interphase when DNA damage or underreplicated DNA is carried over from the previous cell cycle. Like MN-bodies, 53BP1 bodies show persistent DNA damage and a persistent accumulation of a subset of damage response factors within an otherwise normal interphase nucleus. By incompletely understood mechanisms, 53BP1 bodies are proposed to shield DNA lesions until they can be repaired later in the cell cycle. Interestingly, 53BP1 bodies also exhibit transcriptional suppression, again for unclear reasons and by unclear mechanisms^35^. We speculate that a beneficial 53BP1 body response to low-level DNA damage from the previous cell division may cause pathological transcriptional suppression in the setting of the extensive DNA damage and underreplication that occurs on micronuclear chromosomes.

There are two ways in which transcription and chromatin variation from micronuclear chromosomes could be translated into phenotypic variability and long-term epigenetic changes during tumor evolution. One model is that the transcriptional alterations that we document over two generations, can be stably propagated and selected for over many generations. However, our finding that a substantial fraction of micronuclear chromosomes restore transcription after reincorporation into a normal nucleus disfavors this model. Alternatively, the reduction of transcription that we observe could be transient, lasting only a few generations. Transient transcriptional suppression could nevertheless provide considerable phenotypic variation if the prevalence of cells with micronuclei in the population is high, even if the probability of fixation of transcriptional alterations is low. Indeed, a high prevalence of nuclear abnormalities can occur during tumorigenesis because of the interconversion of micronuclei and chromosome bridges, which drives multigenerational cycles of bridging and micronucleation^37^. Therefore, phenotypic effects from ongoing chromosomal instability may be amplified by inherently coupled variation in gene expression.

## Methods

### Cell culture and cell line construction

Cells were cultured at 37ºC in 5% CO2 atmosphere with 100% humidity. Telomerase-immortalized RPE-1 retinal pigment epithelium (CRL-4000, ATCC), U2OS osteosarcoma (HTB-96, ATCC) cells and derivative cell lines were grown in Dulbecco’s modified Eagle’s medium DMEM/F12 (1:1) without phenol red media (Gibco) supplemented with 10% FBS, 100 IU ml^-1^ penicillin, and 100 µg ml^-1^ streptomycin. For cell lines with doxycycline-inducible constructs, tetracycline-free FBS (X&Y Cell Culture) was used.

Stable cells lines H2B–eGFP and TDRFP–NLS RPE-1, mRFP–H2B and eGFP–BAF RPE-1, mRFP–H2B RPE-1 and TDRFP–NLS U2OS were generated by transduction of RPE-1 or U2OS cells using lentivirus or retrovirus vectors carrying the genes of interest as described in^24^.

#### Generation of cells expressing SNAP-MDC1

The RPE-1 GFP-H2B RFP-NLS SNAP-MDC1 cell line (Fig. 3 and Extended Fig. 8) was generated by lentiviral transduction of the SNAP-MDC1-bearing lentiviral vector. This vector was generated by cloning a synthesized SNAPf fragment (sequence from “pBS-TRE-SNAPf-WPRE”; plasmid #104106; Addgene) with AgeI and BstBI restriction sites into the pLenti CMV/TO GFP-MDC1 (779-2) (plasmid #26285, Addgene, gift from E. Campeau; Genewiz) backbone, substituting SNAPf with EGFP at the N-terminus of MDC1. Stably transduced cells were selected by FACS 10 days after transduction for SNAP-MDC1 expression.

#### Generation of the modified U2OS 2-6-3 transcription system

Our modified U2OS 2-6-3 cells contain: GFP-H2B, Cuo-LacI-SNAP, MS2-Halo (Fig. 2, Fig. 4 and Extended Data Fig. 7). These cells were generated from the original U2OS 2-6-3 cells^25^ (gift from D. Spector). Briefly, the “2-6-3” transgene consists of 256 tandem copies of the lac operator enabling visualization of the transgene genomic locus, 96 tetracycline response elements (TREs) to control the reporter transgene, and 24 MS2 translational operators (MS2 repeats) for the visualization of the reporter nascent transcript^25^. The 2-6-3 transgene was introduced into a single euchromatic locus on Chr 1p36^25^. We modified the system as follows. We introduced a lentivirus with the coding sequence of LacI fused to SNAP, under control of a cumate-inducible promoter. Independent control of LacI-SNAP and the MS2 reporter allowed the identification of the reporter in micronuclei in generation 1, followed by assessment of MS2-marked transcription in generation 2. We also stably introduced the genes expressing LacI-SNAP, rtTA, and the MS2 Coat Protein, MCP, used for visualizing the MS2 aptamers.

Specifically, U2OS 2-6-3 were transduced with pLenti CMV rtTA3 Blast (w756-1, plasmid #26429; Addgene, gift from E. Campeau) for the expression of rtTA, a lentiviral vector, phage ubc nls ha 2xmcp HALO, for the expression of “MCP-Halo” (plasmid #64540; Addgene, gift from J. Chao) and lenti Cuo-LacI-SNAP, for the expression of LacI-SNAP. Our LacI-SNAP expression vector, CuO-LacI-NLS-SNAPf, contains the coding sequence for the SNAPf-Tag (sequence from “pBS-TRE-SNAPf-WPRE”; plasmid #104106; Addgene) followed in frame with coding sequence for LacI-NLS (sequence taken from “Cherry-LacRep”; plasmid #18985; Addgene). This sequence was subcloned in the pCDH-EF1-CymR-T2A-Puro (QM200VA-1; System Biosciences SBI), using NheI and BstBI restriction sites. The final modified U2OS 2-6-3 cell line was obtained by selection for hygromycin resistance conferred by the “2-6-3” transgene, for blasticidin resistance, conferred the rtTA expression construct, followed by FACS sorting to identify MCP-Halo and LacI-SNAP expression. Note that LacI-SNAP was transiently induced with 1mM Isopropyl β-D-1-thiogalactopyranoside (IPTG) prior to the FACS sorting. Transient induction was done to avoid genetic instability from LacI binding to the Lac operators, which are a barrier to replication fork progression. The full maps of the constructs used are available upon request. All cell lines used in this study were monitored for mycoplasma contamination.

#### Generation of RPE-1 megaDam cells

RPE-1 3xCherry Dam AID Smash cell line (RPE-1 megaDam) (Fig. 4 and Extended Data Fig. 9), was generated by lentiviral transduction of the megaDam vector into an RPE-1 cell line that has a doxycycline-inducible transgene expressing the E3 ligase, OsTIR1, integrated at the ROSA26 locus^38^. The megaDam vector (Fig. 4a) was generated by synthesizing (Genewiz) a sequence containing three copies of mCherry (based on the sequence from pHAGE-EFS-N22p-3XRFPnls; plasmid #75387; Addgene); the sequences encoding the mAID and SMASh degrons (from^38^). The Dam coding sequence was taken from “TS52_pT_damonly” (a gift from B. van Steensel). The sequence encoding the Dam-mCherry-double degron fusion was cloned and introduced into the lentiviral vector pCW57.1 (plasmid #41393; Addgene, gift from D. Root, Genewiz). A stably expressed RPE-1 megaDam cell line was obtained by antibiotic selection (puromycin).

### Cell cycle synchronization and methods to generate micronuclei

To synchronize cells and generate micronuclei, most experiments in this study used a previously described nocodazole block and release protocol^13,20,24^. Briefly, approximately 15 hours after p53 siRNA (Horizondiscovery) cells were treated with 100 ng/ml nocodazole for 6 hours, followed by a mitotic shake-off procedure. Alternatively (for Fig. 4a-c, Extended Data Figs 4e and Extended Data Figs 9), cells were synchronized at G2/M border by treatment of 9 µM RO-3306 (MilliporeSigma), a CDK1 inhibitor, for 18 hours. G2/M-arrested cells were next released into mitosis by washing five to seven times with medium, followed by addition of 1 µM NMS-P715 (MilliporeSigma) to impair chromosome segregation by inhibition of the MPS1 kinase^39^.

### Nascent EU-marked transcript detection

To detect nascent transcripts, cells were incubated for 30 min with 1 mM 5-ethynyl uridine (EU), which was added approximately 23 hours after mitotic shake-off from nocodazole release. EU incorporation was detected using the Click-iT RNA Alexa Fluor 488 Imaging Kit according to the manufacturer’s instructions (ThermoFischer Scientific).

### Nascent EU-marked transcript detection

The method for targeting a specific chromosome arm to a micronucleus in RPE-1 cells, complete characterization of the editing efficiency of the sgRNA used in this study and the frequency of generation of micronuclei harboring the targeted chromosome are described in detail in^23^. Briefly, the Trueguide Synthetic gRNA system (Thermo Fisher Scientific) was used to generate the sgRNA for chr5q with the sequence G*U*U*GGCCUCCCAAACCACUA (asterisks indicate modified 2’-O-Methyl bases with phosphorothioate linkages). RPE-1 GFP-H2B RFP-NLS cells were synchronized in G0 by serum starvation for 23 hours and were then transfected with the Cas9/gRNA RNP complexes 22 hours post release. Cell synchronization and transfection were performed on cells seeded onto MembraneRing 35 rings (#415190-9142-000; Carl Zeiss), allowing cell isolation by laser capture (see below). Live cell imaging started 3-5 hours post transfection and cells with micronuclei and their siblings were followed until in late G2 phase before cell capture for scRNAseq.

### Cas9 RNP transfection and measurement of editing efficiency

The method for targeting a specific chromosome arm to a micronucleus in RPE-1 cells, complete characterization of the editing efficiency of the sgRNA used in this study and the frequency of generation of micronuclei harboring the targeted chromosome are described in detail in^23^. Briefly, the Trueguide Synthetic gRNA system (Thermo Fisher Scientific) was used to generate the sgRNA for chr5q with the sequence G*U*U*GGCCUCCCAAACCACUA (asterisks indicate modified 2’-O-Methyl bases with phosphorothioate linkages). RPE-1 GFP-H2B RFP-NLS cells were synchronized in G0 by serum starvation for 23 hours and were then transfected with the Cas9/gRNA RNP complexes 22 hours post release. Cell synchronization and transfection were performed on cells seeded onto MembraneRing 35 rings (#415190-9142-000; Carl Zeiss), allowing cell isolation by laser capture (see below). Live cell imaging started 3-5 hours post transfection and cells with micronuclei and their siblings were followed until in late G2 phase before cell capture for scRNAseq.

### Live cell imaging

For the majority of the live-cell imaging experiments, images were collected on Nikon (Ti-E) or (Ti2) widefield inverted microscopes equipped with Perfect Focus, an environmental enclosure to maintain cell culture conditions (37*°*C and humidified 5% CO2), a 20×/0.75 NA Plan Apochromat Lambda objective (Nikon) or a 40×/0.95 NA Plan Apochromat Lambda objective, and a Zyla 4.2 sCMOS camera (Andor). At each timepoint three 2-µm-spaced Z-focal plane image stacks were acquired. Live imaging by confocal microscopy (for the experiments with the adapted U2OS 2-6-3 and MDC1-expressing cells, see below) was performed at 15 min time intervals on a Ti2 inverted microscope fitted with a CSU-W1 spinning disk confocal head (Nikon). At each timepoint three 2-µm-spaced Z-focal plane image stacks were acquired using a 40×/0.95 NA Plan Apochromat Lambda objective. The microscopes were controlled using Metamorph software (Molecular Devices) or NIS-Elements AR (Nikon).

#### Live-cell imaging for Look-Seq2 experiments

Imaging for Look-Seq2 experiments (Fig. 1, Fig. 2 and Extended Data Fig. 1 - Extended Data Fig. 3, Extended Data Fig. 5 and Extended Data Fig. 6) was performed with a widefield inverted microscope and a 20x objective (see above).

#### Live-cell imaging of U2OS 2-6-3 transcription reporter-containing cells

For live-cell imaging of our modified U2OS 2-6-3 nascent transcript reporter cells (Fig. 2, Fig. 4 and Extended Data Fig. 7) cells were seeded on 35-mm ibiTreat Grid-500 dishes (Ibidi) with a gridded imaging surface after mitotic shake-off. SNAP- and Halo-tagged proteins were labeled using 250 nM JF549-cpSNAP-tag and 50 nM JF646-HaloTag ligands (Janelia Materials) for 15 min prior to the start of imaging. To induce the LacI-SNAP expression, cumate (30 ug/ml; System Biosciences) was added in the media immediately after the mitotic shake-off and was washed out prior to imaging. Doxycycline (1 ug/ml; MilliporeSigma) was used to induce the expression of the MS2 transcription reporter approximately 2 hours prior to the start of imaging and was maintained in the medium for the remainder of the experiment (see Fig. 4d). Confocal imaging started 16-19 hours after the mitotic shake-off and was performed as described above for about 24 h or until most of the cells of interest had divided and could be imaged in the generation 2.

#### Live-cell imaging of MDC1-expressing cells

For the live-cell imaging experiments tracking damaged micronuclear chromosomes marked by MDC1 (RPE-1 GFP-H2B RFP-NLS SNAP-MDC1 cells; Fig. 3 and Extended Data Fig. 8), cells were seeded in 35-mm ibiTreat dishes (ibidi) after mitotic shake-off and SNAP-tagged proteins were labeled using 250 nM JF646-SNAP-tag ligand (Janelia Materials) for 15 min prior to the start of imaging. Imaging was started 16-19 hours after the mitotic shake-off and was collected at 15 min intervals with a 40x objective and 2 × 2 image stitching. The exception was the experiments of Fig. 3a, where 35-mm ibi-Treat dishes (ibidi) were used and we collected images at 6 min time intervals without image stitching.

### Capture of single-cells for Look-Seq2

The isolation of live cells for scRNA-Seq was performed in two ways. Our initial experiments were performed in an analogous manner to our early Look-Seq procedure^13^ for single cell whole genome sequencing (which was subsequently refined in^20^). For these experiments, cell isolation was accomplished by trypsinization followed by FACS sorting of single-cells into 384-well µClear imaging plates (Greiner). Micronucleated cells were identified by imaging, and then after the division of these cells, daughters were isolated for scRNA-Seq by trypsinization and replating into 384-well µClear plates after serial dilution. This procedure was employed for a subset of the generation 2 experiments to assess the effect of micronuclear chromosome reincorporation into normal daughter nuclei (10/127 of the total generation 2 samples (see Supplementary Table 1). For all the generation 1 samples and for the majority of the generation 2 samples, we used the laser capture microdissection (LCM) procedure described below (Fig. 1, Fig. 2, Extended Data Fig. 1 - Extended Data Fig. 3, Extended Data Fig. 5 and Extended Data Fig. 6) and and Supplementary Tables 1 and 2). The main advantages of our modified system over previous LCM methods^40^ are (1) minimized cell stress (because the cells are kept in media in the microchamber setup that prevents the culture to dry out) throughout capturing, (2) higher throughput and,(3) an enhanced ability to capture cell families (again because the cells are maintained in media throughout capture of multiple cells).

#### Live-cell imaging for Look-Seq2 experiments

Cells were treated as described in “Cell cycle synchronization and methods to generate micronuclei” for the induction of micronuclei. After mitotic shake-off, cells were handled as described below.

For experiments using the older cell capture method^13,20^, cells were plated into 384-well µClear imaging plates and imaged using widefield fluorescence microscopy at time intervals of 15 min for up to 48 h or until the majority of cells had progressed through mitosis. For LCM capture experiments, cells were instead plated on MembraneRing 35 rings (hereafter “membrane rings”; #415190-9142-000; Carl Zeiss) (see “Single-cell capturing” section) and imaging was performed at 15 min intervals as described above, with the difference that image stitching (2 × 2) was used to track mobile cells across different fields of view.

In many cases we were able to capture the daughter cells (D1 and D2) of the micronucleated mother cell as well as the progeny of the micronculeated cell’s sister (nieces N1 and N2). In some cases, we were unable to capture both niece cells for technical reasons, but these families were included in our analysis if sufficient information could be obtained from the cells captured (e.g., a monosomy in one niece implies a 1:3 segregation pattern even without the biological replicate from the other niece cell, Fig. 2a). We considered micronuclear NE rupture (loss of RFP-NLS) to occur in G1, if rupture occurred within 8 hr of mitotic shake-off and in S/G2, if rupture occurred at any timepoint during interphase after 8 hr. GFP-BAF was used for the assessment of NE integrity in a subset of the “generation 1” samples, as described before^24^.

Development of the modified LCM capture method. We adapted a previously developed LCM system (Palm Microbeam, Carl Zeiss) and re-designed the capturing and imaging setup as described below and in Extended Data Fig. 1b. A custom designed aluminum adapter was constructed using computer numerical control (CNC; SeqTech) milling, in order to be able to perform imaging of cells plated on the membrane rings. We also custom designed and 3D printed an adapter usingVeroWhite material (opaque white Polyjet resin; SeqTech) to allow placement of the membrane rings in a flipped orientation on the Palm Microbeam LCM microscope with a DishHolder 50 CC (#415101-2000-841; Carl Zeiss, see Extended Data Fig.1b). This allowed capturing of cells in a multi-well capture plate that could be placed very close to the cells, which helped increasing the capturing speed, efficiency and thus the throughput of the method. The designs of the adapters are available upon request. The Look-Seq2 method is described in detail as part of the provisional patent with reference number PCT/US20 19/023696, Sep 26, 2019. A hydrophobic barrier was applied at the periphery of the surface of the membrane rings using an immedge pen (Vector Laboratories), in order to prevent media evaporation. Next, the cells were plated on the membrane rings.

At the time of cell capturing, the cells were supplemented with media containing HEPES. Next, the membrane rings were flipped upside down and positioned on the custom-made adapter, after the application of a glass 20 mm glass coverslip (Neuvitro), which we refer to as the “microchamber” (Extended Data Fig. 1b). The cells in the microchamber were transferred to the Palm Zeiss LCM microscope on the custom-made adapter. Cells of interest were identified by extrapolation of the coordinates on the imaging microscope to the LCM microscope using a custom MatLab script and reference marks that were applied to the membrane rings. Snapshots of all the channels of the cells of interest were taken immediately before laser capture microdissection in order to ensure accurate assessment of the micronuclear NE integrity and cell viability (from the maintenance of nuclear RFP-NLS). Cells of interest on small membrane surfaces were then catapulted into single wells in 5.5 ul of lysis buffer (see “Single cell RNA sequencing” section below) in a 96-well capture plate (CapturePlate 96 (D); # 415190-9151-000; Carl Zeiss). The cell lysates were quickly transferred in 96-well PCR plates (Eppendorf) by centrifugation and stored in −20*°*C for cDNA library generation and single cell RNA sequencing.

### Single-cell RNA sequencing

cDNA synthesis and amplification was performed using a modified protocol for the SMART-Seq v4 Ultra Low Input RNA Kit for Sequencing (Takara Bio). Briefly, the manufacturer’s instructions were followed except that 3ul of RNAse Inhibitor were added per 20ul for the “10x Reaction Buffer” solution and all the reaction volumes were decreased by half to maintain reactant stoichiometry. The cDNA amplification from single cells was performed by PCR for 21 cycles and the amplified products were purified using AMPure XP paramagnetic beads (Beckman Coulter).

The quality and quantity of the amplified cDNAs were assessed using the dsDNA HS Assay Kit on a Qubit fluorometer (ThermoFisher Scientific) and Agilent High Sensitivity DNA kit on a 2100 Bioanalyzer system (Agilent Technologies). Samples that were below 0.2 ng/µL and/or had a fragment size distribution that did not match the typical full-length mRNA distribution (with the majority of fragments at around 2 Kb) were excluded. Sequencing libraries were generated by tagmentation using Nextera XT DNA Library Preparation Kit (Illumina) with minor modifications from the manufacturer’s instructions. Briefly, 0.1-0.2 ng/µL of cDNA samples were used in a quarter of the suggested volumes for all subsequent reactions. The Nextera XT Index Kit v2 Sets A-D (Illumina) barcodes were used for the amplified sequencing libraries and the quality of libraries was assessed using a Qubit fluorometer (ThermoFisher Scientific) and a 2100 Bioanalyzer (Agilent Technologies). Sequencing of the libraries was performed either on MiSeq, Hiseq 2500 and NovaSeq 6000 sequencers using mostly paired-end sequencing of 100 bp reads, after quantity normalization and quality assessment of the individual libraries by low-pass sequencing on a MiSeq Nano flow cell.

### Single-cell RNA-Seq data processing and analysis

The complete workflow was implemented as a snakemake pipeline that is publicly available at https://github.com/nikashmyn/scRNA-seq-pipeline.git. Details of individual steps are described below.

#### Alignment and post-alignment processing of sequencing data

Sequencing reads were aligned using STAR 2.7.6a (https://github.com/alexdobin/STAR) to the Gencode v25 reference (–twoPassMode basic; –quantMode: TranscriptomeSAM and GeneCounts) and sorted by genomic coordinate. For post-alignment processing, we followed the GATK’s best practice including adding read group information and executing SplitNCigarReads (both using GATK [v3.8-1-0-gf15c1c3ef]) but skipped duplicate removal as the estimated fractions of duplicated reads is below 5%.

#### Quality assessment of single-cell RNA data

The STAR program outputs various alignment metrics of the RNA-Seq data. We report the following in Table 1: Percentage of unmapped reads, multi-mapped reads, reads mapped to no features according to gene annotations, and reads mapped to multiple features according to gene annotations; Number of genes (transcripts) represented by at least 1, 5, or 10 reads; and the Average number of reads covering each gene. The primary metric for removing poor-quality RNA-Seq libraries is the number of genes covered by ≥5 reads. For control (untreated) RPE-1 cells isolated by FACS, Look-Seq, or Look-Seq2 procedures, we exclude cells with < 6,000 genes covered by five or more reads; for MN related cells, including MN cells, MN sisters, MN daughters, and MN nieces, isolated by either Look-Seq and Look-Seq2, we exclude cells with < 4,000 genes with five or more reads. A total of 619 cells were included in the analysis, 579 of which were sequenced on HiSeq (Illumina) and another 40 by MiSeq (Illumina). All of these samples were listed in Table 1 with annotations of the experimental setup.

#### Quantification of Gene Expression

We calculated TPM (Transcripts per Million) for each gene with RSEM (https://deweylab.github.io/RSEM/) by rsem-calculate-expression.

#### Quantification of allele-specific expression

We generated allelic depths of RNA-Seq reads at heterozygous sites using the UnifiedGenotyper module of GATK [v3.8-1-0-gf15c1c3ef]. The list of heterozygous variants and the haplotype phase of variant genotypes on parental chromosomes were both taken from a prior study^22^. For each gene, the allele fractions of mRNAs from both parental homologs were calculated as the average allelic fraction of variant genotypes in exons and UTR regions.

#### Selection of genes for assessing transcriptional variation due to DNA copy-number variation

Although the average transcription yield of each gene is expected to scale linearly with the gene copy number, the observed mRNA abundance is not strictly proportional to the DNA copy number due to both intrinsic and extrinsic transcriptional noise. To ameliorate such noise and determine the pattern of chromosome segregation both in the cell division giving rise to micronuclei (generation 1) and in the subsequent cell division where the MN chromosome is reincorporated (generation 2), we first identified genes whose transcriptional levels are less affected by transcriptional noise. We excluded genes with very low expression (with fewer than 15% of controls showing TPM values above 5) that display more cell-to-cell variability due to both transcriptional noise^41,42^ and technical variability^43,44^. We further excluded genes with very high expression (top 1%) as these genes may contribute disproportionately to the normalized TPM ratio calculated from the total transcription from a single chromosome or chromosome arm.

#### Quantification of variation in total and allele-specific expression

For each gene, the mean TPM from all control RPE-1 cells (defined in Table 1) was used to normalize the expression in each single cell to calculate normalized expression (TPM ratio). As TPM represents relative transcript abundance, the TPM values in a single cell sample may be uniformly affected by significant changes of highly transcribed genes due to their large contributions to total mRNA content. To suppress such effects and more accurately assess global gene expression variability, we further normalized the TPM ratios in each cell by their mean value in the set of selected genes (see previous section). (The mean of TPM ratios in each cell should be close to 1 if the transcriptome is similar to that of a disomic cell. If a few genes cause a disproportionate change to total mRNA copy number, the change to TPM ratios of the remaining genes is essentially normalized away by this scaling.) We used the normalized TPM ratio to calculate both chromosome (arm)-level and bin-level (50 genes identified in control RPE-1 cells by criteria described in the previous section) average transcription yield to identify changes in the number of transcriptionally active chromosomes or loci.

Due to sparse numbers of heterozygous sites in mRNA reads and because of intrinsic noise of gene transcription from the two homologues, few sites show bi-allelic transcription. We therefore estimate allelic expression based on the number of genes with transcripts derived from either parental homolog, rather than the total number of transcripts phased to each homolog. The aggregated allelic expression is similarly normalized using control RPE-1 cells.

#### Calibration of normal transcriptional variation in disomic, monosomic, and trisomic chromosomes

We first estimated the range of normal transcriptional variation of disomic chromosomes by calculating the mean and standard deviation of average TPM ratios of all chromosomes in control RPE-1 cells. There are two caveats of this calculation. First, as only one chrX copy is actively transcribed (the other is epigenetically silenced by X-chromosome inactivation), the observed transcription levels for Chr. X should reflect transcriptional variation of monosomic chromosomes. Second, as RPE-1 cells all contain an extra copy of Chr.10q (from 60.78Mb to the q-terminus), transcriptional variation in the 10q arm should reflect that of trisomies instead of disomies. We excluded both Chr.X and Chr.10q from the estimation of normal transcriptional variation of disomic chromosomes. We further observed significantly higher variability of the expression of gene-poor chromosomes including Chrs. 18, 21, and 22 in control RPE-1 cells. Due to uncertainties in the inference of the DNA copy number of these chromosomes, we opted to exclude them from the Look-Seq2 analysis.

To estimate the mean and standard deviation of TPM ratio of monosomic or trisomic chromosomes, we took advantage of mis-segregation events that are generated in Look-Seq and Look-Seq2 experiments by nocodazole block and release. Although the average transcriptional yield is expected to be proportional to gene copy number, the average TPM ratio may deviate significantly from the integer DNA copy-number ratio due to transcriptional noise. To identify bona fide trisomies and monosomies, we employed three criteria. First, we required that there need to be approximately proportional changes in the chromosome-wide average TPM ratio (0.5 for monosomies and 1.5 for trisomies). Second, the allele fractions should be consistent with the expected copy-number change (0:1 in monosomies and 1:2 in trisomies). Finally, and most importantly, the monosomies and trisomies need to be either shared between sibling cells (i.e., a pre-existing aneuploidy from a prior cell division) or result from reciprocal changes between sibling cells (i.e.., a de novo mis-segregation occurring in the examined cell division). As sporadic transcriptional variation is unlikely to cause significant chromosome-wide transcriptional amplification or reduction in both sibling cells, the requirement of all three criteria should rigorously identify monosomies or trisomies. Although it is tempting to use clonal trisomic RPE-1 cells as an additional reference (and we have indeed generated single-cell RNA data of trisomic RPE-1 cells), we reasoned that cells from trisomic clones could have acquired long-term adaptive transcriptional changes related to the specific trisomic chromosomes that are distinct from transient transcriptional changes due to sporadic chromosomal copy-number changes. We therefore only used the transcriptional data of spontaneous trisomies as reference. Reference monosomies and trisomies are annotated in Supplementary Table 2 (from both generation 1 and generation 2 samples).

#### Calibration of normal transcriptional variation in disomic, monosomic, and trisomic chromosomes

Based on the average TPM ratios of disomic chromosomes in control RPE-1 cells and monosomic/trisomic chromosomes in nocodazole treated RPE-1 cells, we classified all other chromosomes against these reference distributions using two-tailed z-tests (Extended Data Fig. 1D). For trisomic and monosomic classifications, we used a standard threshold of p ≥ 0.05 (i.e., chromosomes with average TPM ratio within 95% from the reference distribution are classified as the reference); for disomic chromosomes, we employed a Bonferroni correction to account for the presence of 20 disomic chromosomes in each RPE-1 cells (p ≥ .05/20 = .0025). Chromosomes with significant deviations from integer copy-number states were classified as intermediate. The counts of chromosomes classified into each category and the mean and standard deviation of average TPM ratios in both the reference set and the classified set are all shown in Extended Data Fig. 1D. The almost identical ranges of the reference dataset and the newly classified data validates the consistency of this classification.

#### Inference of identity, segregation pattern, and transcriptional yield of micronuclear chromosomes

For each cell in the experimental group, we first used the average TPM ratio for each chromosome to identify chromosomes with transcriptional levels significantly deviating from disomic transcription (summarized in Supplementary Table 2). We further used allelic information to validate the copy number of actively transcribed chromosomes inferred from total TPM ratio and then determined the integer DNA copy-number ratio in family member cells (MN cell and MN sister, or MN daughters and MN nieces) based on the transcriptome data. We then compared the observed copy-number ratio to the expected outcomes of different segregation patterns of the MN chromosome in both MN generation (between MN cell and MN sister cell) and MN reincorporation (between MN daughters) to determine (1) whether the observed integer copy ratio is consistent with the chromosome being in the micronucleus; and (2) the parental haplotype of the MN chromatid. See Extended Data Fig. 2a,b for the description of the workflow of MN cell/MN sister cell analysis and Extended Data Fig. 2c for the workflow of MN daughters/MN nieces analysis.

After identifying MN chromosomes and their parental haplotype, we assessed transcriptional deficiency of the MN chromatid using two measures. For generation 1 samples, if the segregation of the MN chromosome is inferred to be 3:1 (MN cell: MN sister), we calculated the ratio of total transcription between the MN cell and its sister; defective transcription of the MN chromatid is reflected in a significantly lower ratio than 3:1; if the segregation of the MN chromosome is inferred to be 2:2, we calculated the normalized transcriptional yield of the MN haplotype; defective transcription of the MN chromatid is reflected in a normalized transcriptional yield that is significantly lower than 1. Both measures and the final assessment of transcriptional deficiency of MN chromosomes are summarized in Table 2, Tab 2. For generation 2 samples, if the segregation of the MN chromosome is inferred to be 3:1 in the first generation, the DNA copy number ratio between the two MN daughters is expected to be 3:2 and the allele-specific copy ratio of the MN haplotype is 2:1, transcriptional deficiency is classified if the total transcription ratio was significantly below 3:2 and the allele-specific copy ratio was significantly below 2:1. If the segregation of the MN chromosome was inferred to be 2:2 in the first generation, the DNA copy number ratio between the two MN daughters was expected to be 2:1 and the allele-specific copy ratio of the MN haplotype was 1:0; for this scenario, we calculated the normalized transcriptional yield of the MN haplotype in two MN daughters and tested whether the combined transcriptional yield is significantly lower than 1 to assess transcriptional deficiency.

### Same-cell correlative live-fixed imaging

For the same-cell correlative live-fixed experiments, using MDC1-expressing cells (Fig. 3b, c), cells were seeded on 35-mm ibiTreat Grid-500 dishes (Ibidi) with a gridded imaging surface. Live-cell imaging was performed using widefield fluorescence microscope as described in “Live-cell imaging” section. At the end of live-cell imaging, cells were immediately fixed by incubation with MetOH, for 10 min at -20^*°*^C. A snapshot of the last imaging frame including a differential interference contrast (DIC) image was taken to visualize the grids of the coverslip dish. The grid coordinate information and the live-cell images were used to locate the cells of interest after fixation and indirect immunofluorescence imaging. For experiments using the RPE-1 RFP-NLS GFP-H2B (Extended Data Fig. 8a) and “modified U2OS 263” cells (Fig. 4d,e), live-cell imaging was performed as described above. At the end of the live-cell imaging, cells were fixed by incubation with MetOH for 10 min at -20^*°*^C for RPE-1 RFPNLS GFP-H2B or 4% paraformaldehyde for 20 min at RT (modified U2OS 263 cells). Cells of interest were located according to the grid coordinate for subsequent indirect immunofluorescence analysis.

### Indirect immunofluorescence and confocal microscopy of fixed cells

Cells were fixed and prepared for indirect immunofluorescence and confocal microscopy as described previously^20,24^.

Images were acquired on a Nikon Ti-E inverted microscope (Nikon) with a Yokogawa CSU-22 spinning disk confocal head with the Borealis modification, or a Ti2 inverted microscope fitted with a CSU-W1 spinning disk. Z-stacks of 0.4-0.7-µm spacing were collected using a CoolSnap HQ2 CCD camera (Photometrics) or a Zyla 4.2 sCMOS camera (Andor), with a 60×/1.40 NA Plan Apochromat oil immersion objective (Nikon).

The following antibodies were used for indirect immunofluorescence in this study: *γ*H2AX (Ser139) (Millipore #05636-I, 1:400), H3K27Ac (Active Motif #39133, 1:200), MDC1 (Abcam #ab11171, 1:1000), MDC1 (Sigma-Aldrich # M2444, 1:1000), RNA PolII S5 (Millipore #MABE954, 1:400), 53BP1 (Santacruz #22760S, 1:100), H3K27me3 (Thermo Fisher #MA511198, 1:1000), H3K9ac (Cell Signaling #9649S, 1:400), H3K9me2 (Cell Signaling #9753S, 1:400), POM121 (Proteintech 15645-1-AP, 1:200). Staining of Dam-methylated DNA in fixed cells was done with purified GFP-tagged m6A-Tracer protein as described in^45^.

### Image analysis of fixed cell samples

Two image analysis pipelines that were used in this study. To characterize the transcription and chromatin alterations in micronuclei (Fig. 1e-g and Extended Data Fig.4), customized ImageJ/FIJI macros were used as previously described^24^. To characterize the re-incorporated micronuclei (Fig. 3, Fig. 4 and Extended Data Fig. 8, Extended Data Fig. 9), a Python-based analysis pipeline was used, with additional preprocessing procedures performed using ImageJ/FIJI software. Both pipelines overall consist of the following steps: (1) Cells of interest were identified, and their primary nuclei were segmented. (2) Micronuclei or re-incorporated micronuclear chromosomes were identified and segmented. (3) Mean fluorescence intensities (FI) for labeled proteins or DNA were quantified over the segmented regions of interest.

#### Analysis of the transcription and chromatin alterations in micronuclei

Analyses of micronuclei were performed as previously described^24^.

#### Image segmentation and region of interest (ROI) identification

First, the 3-dimension (xyz) images of primary nuclei and micronuclei were segmented using the Li or Otsu thresholding method in ImageJ/FIJI with DNA (Hoechst) signal as input. Second, the nuclear segmentations were further refined using ImageJ/FIJI functions ‘Watershed’ and ‘Erode’ to remove connecting pixels bordering abutting nuclei. Third, nuclear segmentations containing primary nuclei and micronuclei were manually selected as regions of interest (ROIs) using the ImageJ/FIJI functions ‘Wand Tool’. ROIs from one single focal plane where primary nuclear and micronuclear DNA signal are on focus were selected by eye and used for the following quantification.

#### Fluorescence intensities (FI) quantification

The mean FI for labeled proteins or DNA was quantified over the selected ROIs from their corresponding microscope channels. For quantification of nuclear proteins (e.g., RNAP2-Ser5ph, RFP-NLS, Fig. 1e-g and Extended Data Fig. 4), or labelled DNA (Hoechst), the mean FI were calculated for micronuclear ROIs and primary nuclear ROIs, respectively. These mean FI were subtracted by the mean FI of the non-nuclear background to obtain the background subtracted mean FI. The background-subtracted mean FI of MN were divided by the background-subtracted mean FI of the corresponding primary nucleus to obtain the MN/PN mean FI ratios. For quantification of histone modifications including H3K27Ac, H3K9Ac, H3K9me2, H3K27me3 and *γ*H2AX, the MN/PN mean FI ratios of these marks were further divided by the MN/PN mean FI ratio of DNA (Hoechst) to obtain the DNA-normalized FI ratios.

In addition, the background-subtracted mean FI of RNAP2-Ser5ph are also shown as exact FIs without normalizing to the background-subtracted mean FI of the corresponding primary nucleus (Extended Data Fig. 4b).

#### Analysis of generation 2 reincorporated micronuclear chromosomes

Analyses of incorporated micronuclear chromosomes were performed primarily using an automated script written in Python. The code is available upon request.

#### Image segmentation and region of interest (ROI) identification

1. All candidate primary nuclei within the 3-dimensional images were identified and segmented either by the Li thresholding method with DNA (Hoechst) signal as the input, or by the Otsu Li thresholding method with RNA Pol2S5 signal as the input. The nuclear segmentations were further refined using binary mask operations similar to the procedures described above for the micronuclei analysis pipeline.
2. A smaller cropped 3-dimensional image was generated for each segmented primary nucleus object to minimize variability in fluorescence signal across the whole image. Only primary nucleus objects located within the middle 50% of our images were analyzed to minimize the uneven illumination due to the large field of view of the camera (2048 × 2048 pixels). From these cropped 3-dimension images, a single focal plane where the MDC1 or ^m6^A-Tracer (hereafter “m6T”) signal is in focus was selected. This single focal plane was determined as the focal plane with the largest standard deviation (SD) in the FI distribution of all pixels from the MDC1 or m6T channel. The cropped xy images and segmentations for each candidate primary nucleus objects were then analyzed.
3. Primary nuclei that contain potential re-incorporated micronuclear chromosomes were located using the presence of large foci of MDC1 or m6T. To identify MDC1/m6T large foci for each candidate primary nucleus, the FI of all pixels within the corresponding nuclear segmentation were quantified to generate a nuclear FI distribution. Positive pixels were defined if their FI > 2 SD above the mean for the nuclear FI distribution. These positive pixels were subject to a size filter (300 pixels) to remove small noise pixels so that only connected-positive pixels larger than the size filter are kept to generate the final ROIs for MDC1/m6T. Additionally, to increase detection accuracy of m6T foci from cells with a variable m6T expression, candidate nuclei of interest were manually screened using ImageJ/FIJI. The xy coordinates of these candidate nuclei were supplemented as additional inputs for the analysis pipeline and used for locating valid nuclei containing m6T foci according to the above criteria (FI > 2 SD and > 300 pixels) using Python.
4. Segmentations for other objects that are used for the analysis were generated. Specifically, the ROIs for the control region of the primary nuclei were defined by excluding the MDC1/m6T segmentations as well as the nucleoli segmentations from the original nuclear segmentation (see step 1). The nucleoli segmentations were generated using the lower 10% of the primary nuclear FI distribution of RNAP2-Ser5ph. This 10% (percentile) cutoff was determined by comparing to the nucleoli segmentations using Fibrillarin positive signals (which are pixels whose FI > 3 SD above the mean for its total nuclear FI distribution, see Extended Data Figure 8e): the highest overlap with the nucleoli segmentations using Fibrillarin positive signal is achieved by using the lower 10% of the nuclear RNAP2-Ser5ph FI as the cutoff for nucleoli segmentations. ROIs or the *γ*H2AX positive areas were defined by *γ*H2AX positive pixels whose FI > 3 SD above the mean for the nuclear FI distribution. The ROI area occupancy ratio of the *γ*H2AX positive pixels within the m6T foci is used to define different levels of *γ*H2AX in reincorporated m6T micronuclei (Fig. 4c, Extended Data Fig. 9e).
5. A randomized control ROI was segmented by randomly picking a smaller area (at a size similar to the MDC1/m6T ROI) within the primary nuclear ROI generated above for each cell containing a MDC1/m6T foci. The random picking process is performed using our Python-based analysis pipeline.

To validate the identification accuracy of MDC1 and m6T foci identification, the ROI segmentations of a random subset of cells were manually examined. The mis-identification rate of our automated pipeline using random subsets of cells is typically lower than 10%. Additionally, for the m6T dataset after quantification (see below), outliers were also manually examined. The mis-identification rate for the outliers of m6T dataset is 25%. These mis-identified m6T foci (N=27) were mostly m6T positive micronuclei right next to the primary nuclei and were distributed near-evenly at the top and bottom of the measurement distribution. These images were excluded during the analysis.

Fluorescence intensities (FI) quantification. The mean FI of labeled proteins or DNA was quantified over the segmented ROIs above from their corresponding microscope channels. All mean FI were then background subtracted by the corresponding mean FI of the non-nuclear background. The background-subtracted mean FI of MDC1 or m6T ROIs (see step 3 above) and the background-subtracted mean FI of the randomized control ROI (see step 4 above) were normalized to the background-subtracted mean FI of the corresponding primary nuclear ROI (see step 5 above) to obtain the normalized mean FI ratios of labeled proteins or DNA.

For quantification of histone modifications including H3K27Ac, H3K9Ac, H3K9me2, H3K27me3 and *γ*H2AX the normalized mean FI ratios of these marks were further divided by the normalized mean FI ratio of DNA (Hoechst) to obtain the DNA-normalized FI ratios. This controlled for signal enrichment due to chromosome compaction.

#### Analysis of incorporated micronuclear chromosome(s) for the correlative live-fixed imaging

For quantification of RNAP2-Ser5ph and H3K27ac in incorporated micronuclear chromosomes marked by MDC1 foci (Fig. 3b,c), the analysis is performed similarly as described except the following differences: 1. The MN-body segmentations were drawn manually along the MDC1-enriched pixels; 2. the control (or PN) segmentations were defined manually as a large PN region excluding nucleoli. ROIs for MN-body and control were manually selected over these segmentations for a single Z plane where MN bodies were on focus. The mean FI MN-body/control ratios were obtained by dividing the background-subtracted mean FI of the MN-body ROIs to the background-subtracted mean FI of the control ROIs.

### Analysis of live-cell imaging data for MS2-marked nascent transcription

To quantify the MS2-based transcription level (Fig. 2e,f, Fig. 4e and Extended Data Fig. 7), an automated script written in Python was used, with assists using ImageJ/FIJI. The code is available upon request.

#### Image segmentation and LacO/MS2 foci tracking

To locate the object of interest, the image series were divided into three parts: generation 1 interphase, mitosis, and generation 2 interphase.

For timeframes covering the generation 1 interphase, all primary nuclei were segmented using the cellpose package^46^, and all LacO/MS2 foci were segmented using the Yen segmentation method^47^ for each timeframe. The primary nuclei and LacI foci of interest in the first timepoints were identified by finding the object segmentation whose distance is minimal to a user-provided xy centroid coordinate of the nuclei and the LacO/MS2 foci, respectively. For the following timepoints, the same nuclei and LacO/MS2 foci were tracked automatically by finding the object segmentation whose distance is minimal to the identified nuclei and the LacO/MS2 foci segmentations from the prior timepoint (or timepoints). Additionally, the distance between the LacO/MS2 foci and their corresponding primary nuclei is also evaluated and considered to assist the tracking of the correct LacO/MS2 foci. The xy centroid coordinates of the segmentations were used for estimating the object moving distance above.

For timepoints around mitosis, the nuclei and LacO/MS2 foci were tracked manually in ImageJ/FIJI to accurately identify the partitioning of LacO/MS2 foci into daughter cells during mitotic exit.

For the interphase of generation 2, daughter primary nuclei were segmented and tracked similarly as described for the first interphase. For LacO/MS2 foci segmentation tracking, we used a combination of several criteria for technical reasons. As long-term binding of LacI to LacO can impair DNA replication, we terminated LacI gene expression at 18 hours post mitotic shake-off. This led to a loss of LacI signal in a subset of daughter cells during the generation 2 interphase, particularly evident at later timepoints. For these cells that had lost the LacI signal, we quantified the FI distribution for the nuclear MS2 signal and identified positive MS2 pixels whose FI > 3 SD above the mean for the nuclear MS2 FI distribution. The enrichment of such nuclear MS2 positive foci (if present) was then used for the LacO/MS2 foci segmentation tracking for the subsequent timepoints. For timepoints where LacI foci persisted, we tracked the LacO/MS2 foci as described for the generation 1 interphase. If no LacO/MS2 foci could be identified during the generation 2 interphase, timepoints were annotated as having no MS2 expression, and therefore no segmentation was performed.

Additionally, we manually examined the nuclei and LacO/MS2 foci tracking because the estimation of object movement using minimal centroid moving distance could lead to incorrect tracking when objects swap positions between timepoints. For these timepoints, additional xy pixel centroid coordinates for the nuclei and LacO or MS2 foci were obtained using ImageJ/FIJI and supplemented the automated object tracking.

#### Fluorescence intensity (FI) quantification for region of interest (ROI)

ROIs for the LacO or MS2 foci and the corresponding primary nucleus were selected from their segmentation, as described above. ROIs from one single focal plane where the LacI signal is on focus were used for FI measurements. Based on these ROIs, the mean FI of the MS2 signal for LacO/MS2 foci and primary nucleus pairs, along with the non-nuclear background was quantified. The background-subtracted mean FI of LacO/MS2 foci was divided by the background-subtracted mean FI of the primary nuclear areas (excluding the LacO/MS2 foci) to obtain the normalized MS2 level.

For timepoints that were annotated as having no MS2 expression, a value of 1.7 was assigned because this value is the minimal detectable MS2 signal above the background. To obtain this value, we analyzed 23 control cells over two cell-cycles whose LacO/MS2 focus was located within the primary nucleus for both generation 1 and 2. We quantified the normalized MS2 level during the generation 2 interphase for timepoints where the cells had lost LacI signal after we stopped the LacI expression. These control cells maintained the MS2 reporter transcription and their LacO/MS2 foci were detected and segmented from MS2 positive pixels (FI > 3 SD above the mean for the nuclear MS2 FI distribution, as described above). The minimum of the normalized mean FI for all (n = 477) detected MS2 positive foci above was 1.7, defining 1.7 as the minimum detectable mean MS2 signal in these experiments. Therefore, 1.7 was used as the normalized MS2 signal when no positive MS2 foci could be detected, which is a conservative estimate which should underestimate the degree of M2S signal loss for micronuclear chromosomes that are in a normal generation 2 daughter nucleus.

### SDS–PAGE and western blotting

Lysis of “RPE1Dam” and control RPE-1 cells (Exended Data Fig. 9c) was performed after trypsinization and washes with PBS by adding an equal volume of a 2× lysis buffer (100 mM TrisHCl pH 6.8, 4% SDS, 12% β-mercaptoethanol). Whole-cell lysates were denatured at 100^*°*^C for 10 minutes, Laemmli-SDS sample buffer (Boston BioProducts) was added, and the samples were subjected to SDS–polyacrylamide gel electrophoresis on NuPAGE 4–12% Bis-Tris gradient gels (Novex Life Technologies). The proteins were then transferred onto a nitrocellulose membrane (Millipore). The membranes were blocked using Odyssey Blocking Buffer (LI-COR) and were incubated with primary antibodies for 1 hour at room temperature or overnight at 4^*°*^C. The primary antibodies and dilutions used were anti-mCherry rabbit 1:1000 (ab167453, Abcam) and anti-GAPDH mouse 1:5000 (ab9485, Abcam). After washes with PBS-T, we incubated the membranes with the fluorescent secondary antibodies IRDye 680RD Donkey anti-rabbit 1:5000 (926-68073, LICOR Biosciences) and IRDye 800CW Donkey anti-mouse 1:5000 (926-32212, LICOR Biosciences), for 1 hour at room temperature. Membranes were visualized using a ChemiDoc MP Imaging System (BioRad). Note that the images shown in Extended Data Fig. 9c are cropped to show the bands at the protein size.

### Fluorescence activated cell sorting

RPE-1 megaDam cells (see “Cell culture and cell line construction” section) were analyzed by FACS for mCherry expression using a LSR Fortessa Flow Cytometer (BD)(Extended Data Fig. 9b). Cells were stained with DAPI for dead cell exclusion and live cells were analyzed for their percentage of mCherry positive cells (excluding auto-fluorescent cells by gating PE relative to FITC). Data were analyzed using FlowJo software (BD).

## Data availability

The authors declare that the data supporting the findings of this study are available within the paper and its Supplemental Information files. Sequencing data will be deposited to the Sequencing Read Archive (SRA). All other data sets generated in this study are available from the corresponding author upon reasonable request.

## Code availability

Scripts and pipelines used for scRNAseq data analysis and for image analyses performed in Extended Data will be available at github on-line repository.

## ACKNOWLEDGEMENTS

We are grateful to Janelia Research Campus, D. Spector, and T. van Schaik for reagents, R. Nicol, H. Zhang and Y. Brody for assistance and advice with laser capture microscopy, Jinyu Wang, Ramya Parasuram, Mitchel Leibowitz and Hawa Ndiaye for help with preliminary experiments, Natalia Sebryn and Rachel Davidowitz for help with scheme illustrations and/or video editing, S. Armstrong for use of an LSR Fortessa Flow Cytometer, and R. Jaenisch, M. Meyerson, A. Spektor and members of the Pellman laboratory for discussions. We thank the Center for Cancer Genomics of Dana Farber Cancer Institute for sequencing services. D.P. is an HHMI Investigator, a member of the HMS/Boston Ludwig Center and is supported by NIH R01 CA213404-24 and an award from the G. Harold and Leila Y. Mathers Foundation.

## Author contributions

S.P. and D.P. conceived the project, S.P. designed the experiments with the supervision of D.P., S.P. performed most of the experiments, S.P. invented the modified LCM system, E.S. helped with experiments, S.P., N.M., E.J. and C-Z.Z. designed the bioinformatic analysis and N.M., E.J. and C-Z.Z. performed the analysis of the scRNAseq data, S.L. developed the automated image analysis pipelines, S.P., N.M., S.L., E.J., E.S., and C-Z.Z. analyzed data, BvS contributed reagents and advise for the DamMN experiments, S.P., C-Z.Z and D.P. wrote the manuscript with edits from all the authors, D.P. supervised the study.

## Reporting Summary

Further information on research design is available in the Nature Research Reporting Summary linked to this article.

## Supplemental Information

**Supplementary Table 1. Summary of single cells for single-cell RNA-Seq analysis**

Cells for which RNA-Seq data have been generated are grouped by the experimental design (Column A) and the identity of the ancestor cell (“Family ID”). Each family consists of cells descended from a single ancestor as identified by live cell imaging and each family member is assigned a unique ID (“Cell ID”). The remaining columns summarize various quality metrics of each single-cell library that are generated by STAR. Control RPE-1 cells (untreated, FACS, Look-Seq2(old capture), and Look-Seq2(LCM) are included if they have > 6,000 genes with five or more reads; MN related cells (Look-Seq2(old capture) and Look-Seq2(LCM), Generation 1 and Generation 2) are included if they have > 4,000 genes with five or more reads.

**Supplementary Table 2. Summary of single-cell analysis of chromosomes with significant deviation from disomic transcription determined using control RPE-1 cells**

*Tab 1: Summary of defective or normal MN transcription assessed from single-cell data of MN cell/MN sister pairs (generation 1, Tab 2) and of MN daughters/MN nieces (Generation 2, Tab 3)*.

*Tab 2: Summary of Look-Seq2 analysis of MN cells and MN sisters (generation 1)*. MN cells (cell ID in column C) and sisters (cell ID in column D) from each family (family ID, column B) are grouped by the status of the NE integrity of the micronucleus (column A) and the presence of micronucleus determined by live-cell imaging. Shown for each family are chromosomes with non-disomic transcription in at least one cell (either MN cell or MN sister). Chromosomes with near normal (disomic) transcription based on the range of transcriptional variation in normal RPE-1 cells are omitted. In Sample F206, all chromosomes display near disomic transcription. We conclude that the MN chromosome underwent 2:2 segregation and was normally transcribed in the MN cell (F206.2), although the identity of the MN chromosome cannot be determined. Columns F-I show the average TPM ratio of the MN cell (Column F), the integer copy number of actively transcribed chromosomes (Column G) inferred based on reference transcription data of disomic, monosomic, and trisomic chromosomes (Extended Data Fig. 1D and Methods), and the allelic fractions of transcripts from both parental haplotypes (Column H and I). Columns J-M display the same data types as in Columns F-I for the MN sister cell. Based on the inferred DNA copy-number ratio between the MN cell (Column G) and the MN sister (Column K) and allelic expression data (Columns H,I,L,M), we determine the integer copy ratio of actively transcribed chromosomes between the MN cell and MN sister (Column N). Chromosomes that were initially classified as having non-disomic transcription (intermediate) based on total gene expression but subsequently inferred to be disomic based on allelic expression are shaded in grey. By comparing the RNA-derived integer copy ratio with expected outcomes from either 1:3 or 2:2 segregation (Fig. 1 and Extended Data Fig. 2), we determine whether the chromosome is MN related (Column O) and the corresponding DNA segregation pattern (Column P). Based on the RNA-derived copy-number ratio (Column N) and the inferred DNA segregation pattern (Column P), we classify the (binary) status of transcription of the MN chromosome (Column Q) and also determine the parental haplotype of the MN chromatid (Column R). If the allelic imbalance between the MN haplotype and the normally segregated haplotype is only visible in one arm, the event is marked as Arm-level (Column S). Columns T-V summarize the main results of the transcriptional analysis of each MN chromosome. For MN chromosomes that have undergone 1:3 missegregation (between MN sister and MN cell) and display lower than expected transcription (“defective”), the ratio of transcription between the MN cell and the MN sister is shown (Column U). For MN chromosomes that have undergone 2:2 segregation and display lower than expected transcription (“defective”), the transcription yield of the MN chromosome (Column V) is directly calculated by multiplying the total transcription yield of the MN sister by the allelic fraction of the MN haplotype. We choose a few monosomic and trisomic chromosomes as reference monosomies/trisomies. These are verified both by the chromosome segregation pattern and by the ratios of total and allelic transcription of MN cells/MN sisters and are annotated in Column W. For two chromosomes (Chr.15 in F230 and Chr.1 in F73, colored blue), we inferred an abnormal integer copy ratio (2:3) between the MN cell and the MN sister cell that is most consistent with a pre-existing trisomy in the mother cell undergoing 3:3 segregation. We note that acrocentric chromosomes (Chr.13-15, Chr.21 and 22 are excluded from the analysis) often display higher transcriptional variability, causing higher uncertainty in the assessment of the integer copy number of actively transcribed chromosomes. In two cases (colored in red) the integer copy ratios assessed from RNA-Seq data (F79:Chr.15 and F76:Chr.13) are difficult to interpret based on the expected segregation patterns. These two chromosomes are excluded from the final summary.

*Tab 3: Summary of Look-Seq analysis of MN daughters and Look-Seq2 analysis of both MN daughters and MN nieces (generation 2)*. Columns A-S are organized similarly as in Tab 2 but include transcription data for both MN daughters and MN nieces (when available from Look-Seq2). As these are from cells that have reincorporated MN (i.e., with no detectable MN), we only have knowledge of the identity of daughters of the MN cell (“MN daughters”) and daughters of the MN sister cell (“MN nieces”); this information is reflected in the column headers. The integer copy ratios of MN daughters and MN nieces (when available from Look-Seq2) are listed in Column T; the inferred DNA segregation patterns in the first generation (between MN cell and MN sister) and in the second generation (between MN daughter A and MN daughter B) are listed in Columns U and V. Chromosomes that were initially classified as having non-disomic transcription (intermediate) based on total gene expression but subsequently inferred to be disomic based on allelic expression are shaded in grey. In one case of Chr.13 and three cases of Chr.19, an initially classified trisomy is re-classified as disomy based on allelic expression; these cases are colored in red. Chromosomes with transcriptional patterns consistent with those due to MN segregation/reincorporation are marked in Column W and annotated in Column X (haplotype) and Y (if only a chromosome arm was affected). We further calculated the allele-specific expression of the MN haplotype in both MN daughters (Columns Z, AA, AB, AC) and either the ratio of allele-specific transcription in Column AD (1:3 missegregation in generation 1, 3:2 segregation in generation 2) or the combined allele-specific transcription in Column AE (2:2 segregation in generation 1, 2:1 segregation in generation 2). The final conclusion about whether transcription of the reincorporated micronuclear chromosome is “defective” or “normal” is summarized in Column AF. Three families which we inferred to have pre-existing trisomy 12 are colored in light blue.

**Supplementary Video 1. Example of a Look-Seq2 experiment**.

The video starts during generation 1. An MN cell (GFP-H2B channel) and its sister are followed. The micronucleus ruptures (loss of RFP-NLS) during the approximate time of S/G2 during generation 1. Both cells divide, generating MN daughters and MN cell nieces in generation 2. The chromosome from the micronucleus is reincorporated into one of the daughters.

**Supplementary Video 2. Live imaging of nascent transcripts after two chromosomes from micronuclei are reincorporated into daughter nuclei**.

The reporter locus is visualized with LacI-SNAP; nascent transcription is visualized with MCP-Halo; and the loss of the general nuclear pool of MCP-Halo indicates rupture of the NE of both micronuclei. One chromosome recovers transcription whereas the other does not.

**Supplementary Video 3. Formation of an MN-body marked by SNAP-MDC1**.

In generation 1, a micronucleus forms and undergoes NE rupture (loss of RFP-NLS). The damaged chromosome from the micronucleus is only decorated with SNAP-MDC1 during mitosis. In generation 2, the SNAP-MDC1 marked chromosome is reincorporated into a daughter nucleus, forming a long-lived MN-body.

---

**Extended Data Figure 1.**
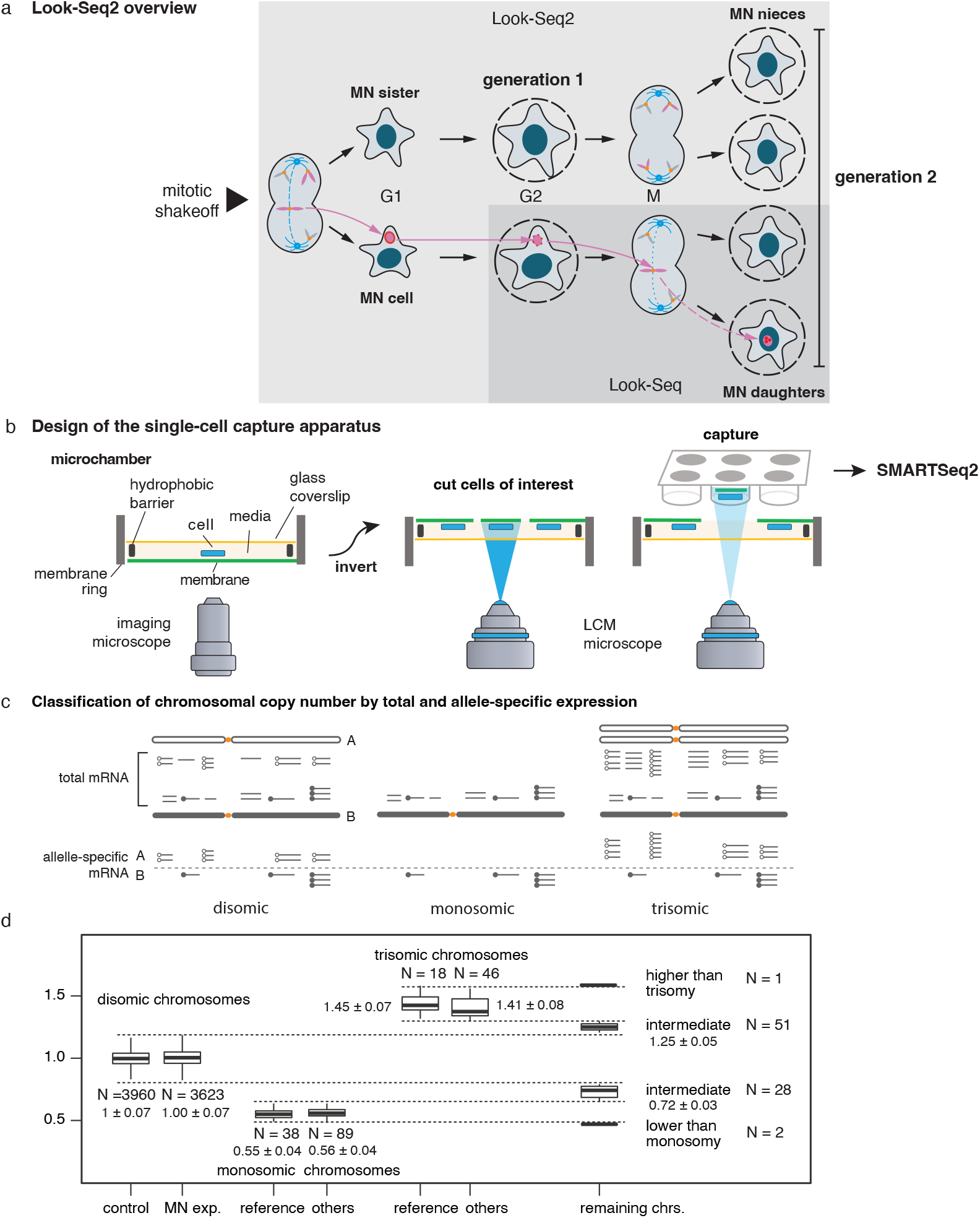
Experimental and Analytical workflows. (a) Scheme of Look-Seq2. There are two key improvements compared to the original Look-Seq. First, live-cell imaging starts before the first cell division that leads to micronuclei; this enables tracking, isolation, and transcriptome analysis of both the MN cell and its sister cell (Fig. 1, “generation 1”). Alternatively, we monitored cells over two cell divisions (Fig. 2, “generation 2”) and identified examples where no micronucleus was detectable after the second division (reflecting reincorporation of the MN chromosome into one or both daughter nuclei). We then analyzed the two nieces of the MN cell and the two daughters of the MN cell. The second improvement is that single cells are isolated using a new capture strategy with minimal mechanical perturbation that is illustrated in b. (b) Second generation experimental strategy for single-cell capture and sequencing. We adapted a previously developed LCM system (Palm Microbeam, Carl Zeiss) and re-designed the imaging and capture setup. The modifications enable the inversion of the membrane rings relative to the microscope objective. This allows medium to be present continuously throughout capture, which provides more time for the capture of family member cells. The set-up is also compatible with laser catapulting into 96 well plates, which further increases throughput. See Methods for details. (c) Two measures of transcription yield from single-cell RNA-Seq data: (1) The abundance of transcripts from all DNA copies of each gene is measured using transcripts per million (TPM) normalized to the mean TPM in diploid RPE-1 cells. The normalized TPM ratio is averaged both across entire chromosomes and in bins of 50 genes to derive estimates of the chromosomal DNA copy number and local transcriptional yield (i.e., the average number of transcriptionally active gene copies). (2) The fraction of transcripts derived from each parental homolog is estimated directly from the counts of haplotype-specific sequencing reads. Details of the computational analysis are provided in Methods. (d) Summary of average TPM ratio in disomic, monosomic, trisomic, and other transcriptional states in: control, unmanipulated RPE-1 cells; the MN experimental set combining all chromosomes; the reference data set of handpicked monosomies and trisomies described above; other classified monosomies or trisomies (e.g, spontaneous trisomy of Chr.12 that is common in RPE-1 cells); and rare samples that exhibited transcription yields greater than trisomy (e.g., rare tetrasomies), intermediate yields (including those due to arm-level copy number alterations), or samples below monosomy (e.g. from rare biallelic loss or loss of active X). Details of the classification procedure are summarized in Methods.

**Extended Data Figure 2.**
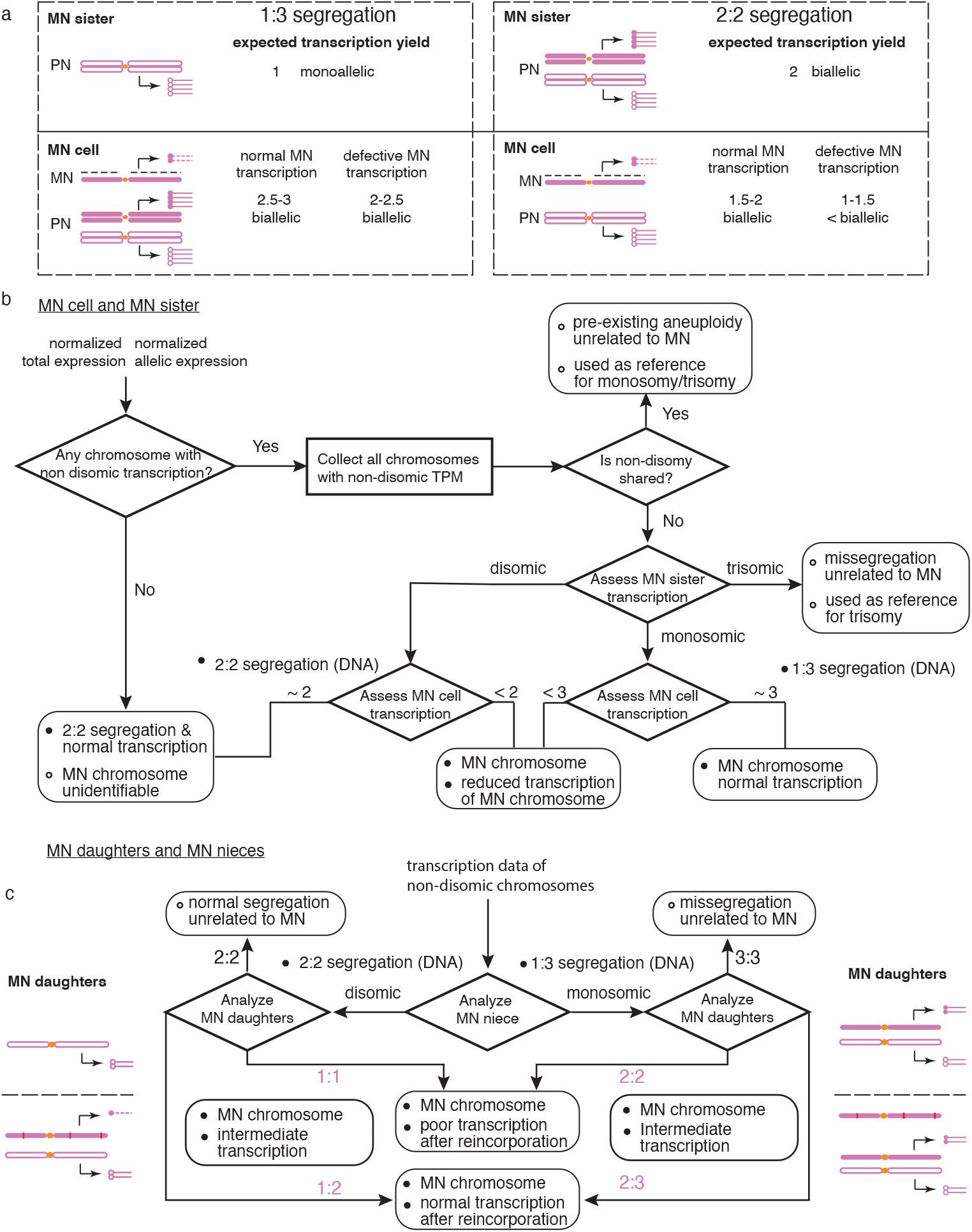
Flowchart of the inference of micronuclear chromosomes, MN chromosome segregation and MN reincorporation from single-cell RNA data. (a) Expected transcription yield of the micronuclear chromosome and its homologous chromosome in the MN sister and the MN cell under 1:3 segregation (left) and 2:2 segregation (right). The representation of the MN chromatids and the intact chromatids follows the same convention as in Fig. 1. For the MN sister cell, it is expected to show monosomic transcription of the intact chromatid (open magenta) in the primary nucleus under 1:3 segregation and bi-allelic disomic transcription of two intact chromatids (open and filled magenta) under 2:2 segregation. For the MN cell, the expected transcription yield varies depending on the transcription yield of the MN chromatid. As the MN chromatid is poorly replicated, its average transcription yield in G2 may be lower than the average transcription yield of the normally segregated homolog even under normal transcription. The expected range of total transcription in the MN cell assumes that the average transcription yield of the MN chromosome if it has normal transcription is at least 50% of that of the normally segregated chromosome in the primary nucleus in G2. (b) Workflow illustrating the logic behind the determination of (1) shared, pre-existing chromosome copy-number changes between the MN cell and its sister cell; (2) chromosome mis-segregation between the MN cell and its sister cell that did not result in MN; and (3) chromosome missegregation events that result in MN. We identify MN-related chromosomes based on the constraint that the only possible transcription ratios between MN cells and MN sisters are 2 + x : 1 (3:1 segregation) or 1 + x : 2 (2:2 segregation, x represents the average transcription yield of the MN chromatid). Thick outlined boxes indicate calculations or logical operations; rounded rectangles indicate conclusions. Note that this analysis requires knowledge of which cell is the MN cell and which cell is the MN sister, which is known from live-cell imaging prior to single-cell sequencing (the advantage of Look-Seq2). Moreover, the identification of MN chromosomes utilizes the asymmetry between MN sisters and MN cells but does not make assumptions about the expected transcription yield of the MN chromosome (x) in the MN cell. The only exception is when the MN chromosome undergoes 2:2 segregation and then displays complete normal transcription (x = 1): In this scenario, the MN chromosome is equivalent to a normal chromosome although we will not be able to determine its identity from the transcriptome data. (c) Inference of MN chromosomes, MN chromosome segregation pattern, and MN chromosome reincorporation pattern from single-cell RNA data. Similar to the inference in (a), we start by assessing the transcription in MN nieces to determine the segregation pattern in the first generation. If the MN nieces show monosomic transcription but the MN daughters show bi-allelic disomic or trisomic transcription, then this pattern indicates a 1:3 segregation in the first generation (Extended Data Fig. 5b and Extended Data Fig. 6b). If the MN nieces show disomic transcription, then either one MN daughter should show monosomic transcription (missing the MN chromatid, Extended Data Fig. 5c and Extended Data Fig. 6a), or both MN daughters show monosomic transcription (one missing the MN chromatid, the other containing a transcriptionally silenced reincorporated MN chromatid), or both MN daughters show intermediate transcription reflecting partial reincorporation of fragmented DNA of the MN chromatid (Extended Data Fig. 6c).

**Extended Data Figure 3.**
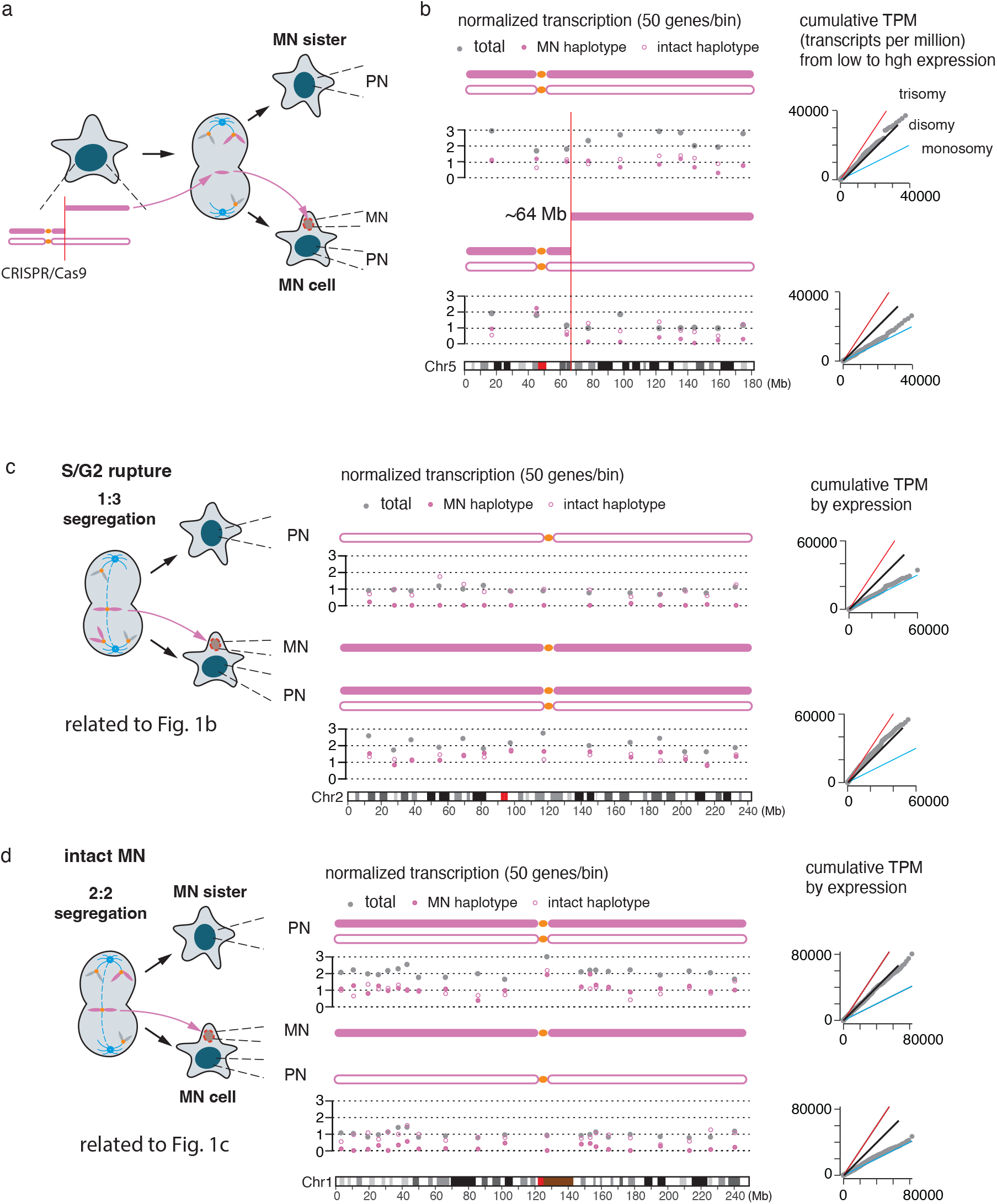
Additional data on the loss of transcription in newly generated MN. (a)The experimental strategy to form MN containing acentric chromosome fragments generated by CRISPR/Cas9 as reported in our recent study ^23^. Because the broken chromosomes can be generated from either one or both sister chromatids from either homolog, there are multiple possible outcomes of DNA segregation (see ^23^); the example shown here demonstrates the capability to identify one outcome when the micronuclei only contain one copy of a broken chromosome arm from single-cell RNA-Seq data. This outcome most closely resembles the segregation patterns generated by nocodazole block-and-release. (b) One possible outcome for the distribution of broken (filled magenta) and intact (open magenta) chromatids to both primary nuclei and the micronucleus, which was corroborated from the transcriptomes of the MN sister cell and the MN cell. Three measures of gene transcription are shown: Filled gray circles are the normalized total TPM ratio; filled and open magenta circles are the normalized allelic expression of the broken and the intact homolog that is determined using the parental haplotypes. All three signals are averaged over bins containing 50 genes expressed in normal RPE-1 cells. The decrease in both total expression and the haplotype-specific expression of the broken chromatid around 64 Mb on 5q indicates that the acentric fragment from 64 Mb to the q-terminus was partitioned into the micronucleus after Cas9-breaks generated at 64 Mb. Reduced transcription of Chr.5 in the MN cell is also evident from the reduction in total transcription relative to normal disomic Chr.5 (black line), as shown in the cumulative TPM plot on the right. As the cumulative TPM plot is generated for all genes on Chr.5, it does not distinguish chromosome-wide transcriptional reduction from regional loss of transcription. (c) Full data analysis for the data shown in Fig. 1b (F216), including the 50-gene averaged yield of total (gray) and allele-specific (open and filled circles) expression, and cumulative TPM (from low to highly expressed genes) plots that validate the inference of monosomic and disomic transcription. The inferred chromosomal compositions in both cells are shown above the bin-level TPM data. (d) Same as (c), above, for Fig. 1c (F220).

**Extended Data Figure 4.**
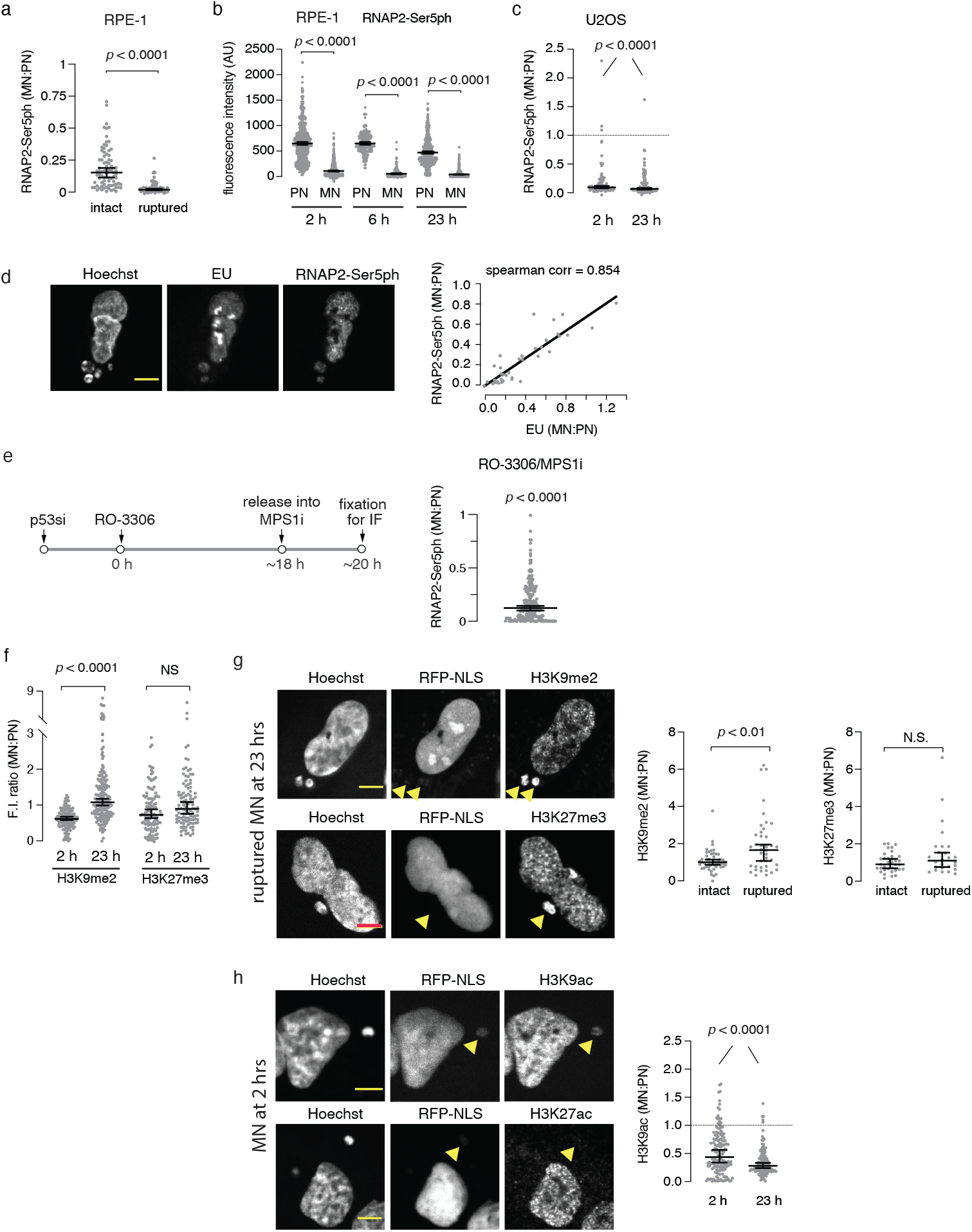
Transcription and chromatin modifications from newly formed (generation1) micronuclei. (a) Transcription is present at a reduced level in intact MN and is nearly absent in ruptured MN. MN:PN fluorescence intensity (FI) ratios are shown for RNAP2-Ser5ph 23 hr post mitotic shake-off. RFP-NLS levels were used to assign the micronuclei in the two groups (n = 83 and 82, left to right, from four experiments). Micronuclei with NLS ratios below 0.1 relative to the PN were considered ruptured and above 0.3 were considered intact. Median with 95% confidence interval (CI); Two-tailed Mann–Whitney test. (b) Data from Fig. 1e, but instead of the MN:PN FI ratios what is shown here is the background normalized intensity values at 2, 6 and 23 h post release from nocodazole and mitotic shake-off (n = 647 for 2 h, 213 for 6 h and 523 for 23 h, from at least two experiments). Median with 95% CI; Kruskal-Wallis with Dunn’s multiple comparisons test. (c) In U2OS cells, active transcription (RNAP2-Ser5ph) is also reduced in newly formed MN. Performed and analyzed as in (a) above (n = 88 and 104, left to right, from two experiments). (d) Independent confirmation of the MN transcription defect by 30 min EU pulse labeling. Left, representative images of cells with S/G2, generation 1 MN. Right, Strong correlation between RNAP2-Ser5ph and EU levels. Cells with varying levels of RNAP2-Ser5ph intensity were selected and then EU intensity levels were measured (n = 37, from one experiment). Two-tailed Spearman’s correlation. Scale bar 5 µm. (e) MN transcription defects are evident in MN generated by an alternative method (G2 arrest with CDK1 inhibition, followed by release into an MPS1 inhibitor). This synchronization and MN induction method differs from the nocodazole block and release protocol primarily used in this study because it shortens rather than lengthens mitosis (excluding hypothetical artifacts from prolonged mitotic arrest). Left: scheme of the experiment. RPE-1 cells were analyzed 2 hours after release from the G2 block (n = 334, from three experiments). Right: quantification and analysis of the results as in (a), above. (f) Modest increase in repressive chromatin marks in a subset of late S/G2 MN (23 hr post-mitosis). Performed and analyzed as in (a) above (n = 129, 179, 114 and 105, left to right, from two experiments). (g) Left, representative images from (f) of cells with ruptured MN and enrichment for H3K9me2 and HK27me3 in the MN. Arrowheads: micronuclei lacking normal RFP-NLS accumulation. Right: related to (f), but comparing intact and ruptured MN for H3K9me2 (n = 48 and 40, left to right, from two experiments) and H3K27me3 (n = 35 and 26, left to right, from two experiments) at 23 h post mitotic shake-off. Performed and analyzed as in (a) above. Scale bars 5 µm. (h) Loss of H3K9ac and H3K27ac in MN at the indicated timepoint during interphase. Left: representative images of the indicated cells at 2 hours post mitotic shake-off. Right: quantification and analysis of the results for H3K9ac as in Fig. 1g (n = 148 and 124, left to right, from two experiments). Scale bars 5 µm.

**Extended Data Figure 5.**
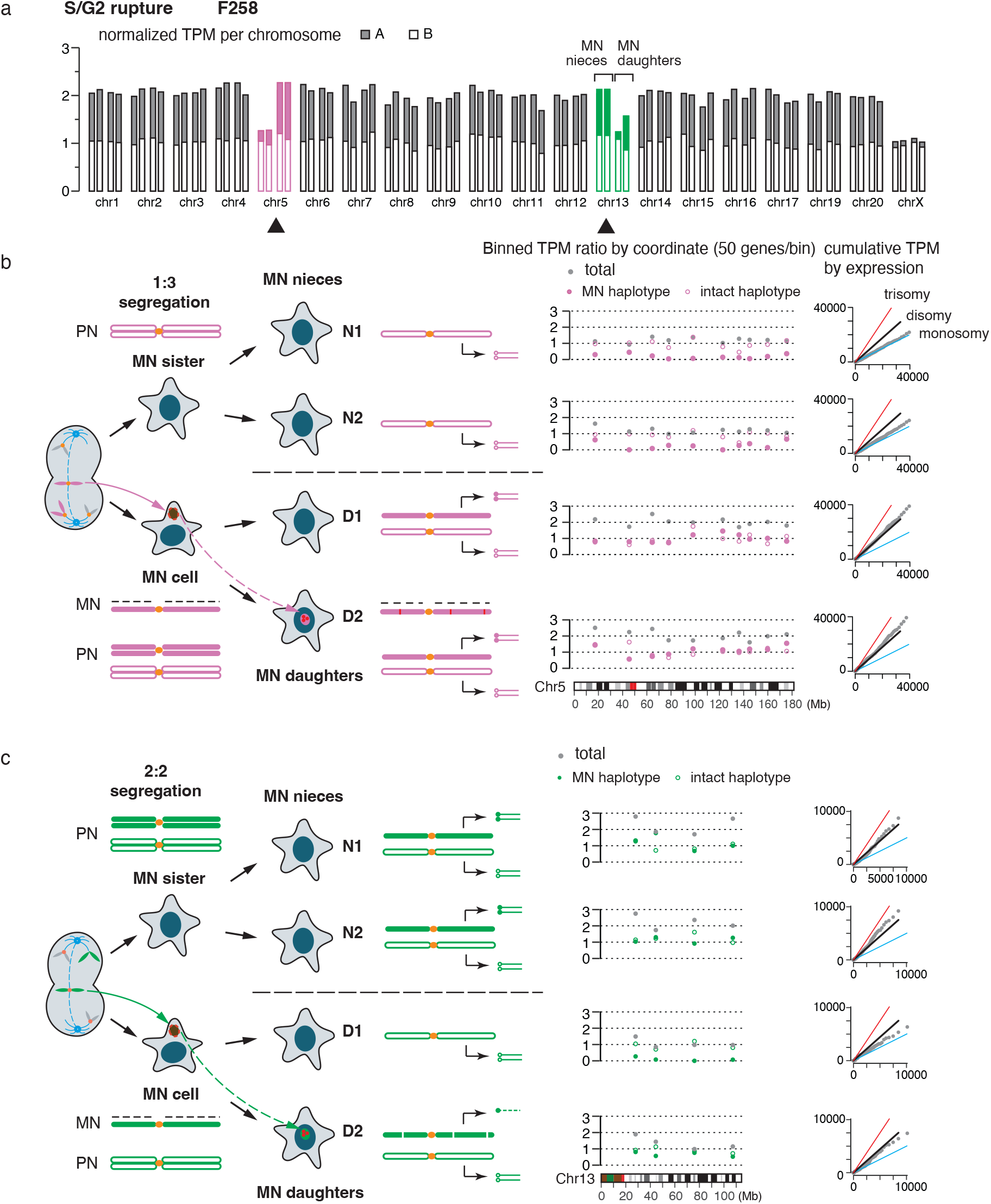
Complete data set for the F258 sample shown in Figure 2b. (a) Average TPM ratio and allelic ratio of all chromosomes (except the low gene density chromosomes 18, 21, 22). Non-disomic transcription of Chr.5 is shown in magenta and Chr.13 is shown in green. The first two cells in the set of four are the MN nieces; the second two cells are MN daughters. Transcription data of Chr.5 and Chr.13 is shown in b and c. (b) The segregation pattern, expected transcriptional yield, and observed transcriptional data of Chr.5 in all four cells. The presence of monosomic expression in both nieces and disomic/biallelic expression in both MN daughters indicate a 1:3 segregation of Chr.5. As the two MN daughters both display close to disomic transcription but one or both of them have reincorporated the Chr.5 copy from the micronucleus, we conclude that the reincorporated Chr.5 is not actively transcribed. (c) The segregation pattern, expected transcriptional yield, and observed transcriptional data of Chr.13 in all four cells. In contrast to the pattern of Chr.5, the two nieces both display disomic/biallelic expression and one MN daughter displays monosomic expression; this pattern establishes a 2:2 segregation of Chr.13. The presence of transcripts phased to the MN haplotype (filled green circles in the bottom cell) indicates transcription of the reincorporated Chr.13 in the bottom cell. However, the total transcriptional yield from both Chr.13 copies is significantly lower than the range of normal disomic transcription, suggesting only partial recovery of the reincorporated Chr.13. We note that Chr.13 contains fewer genes and there is more variation in the transcriptional yield than Chr.5.

**Extended Data Figure 6.**
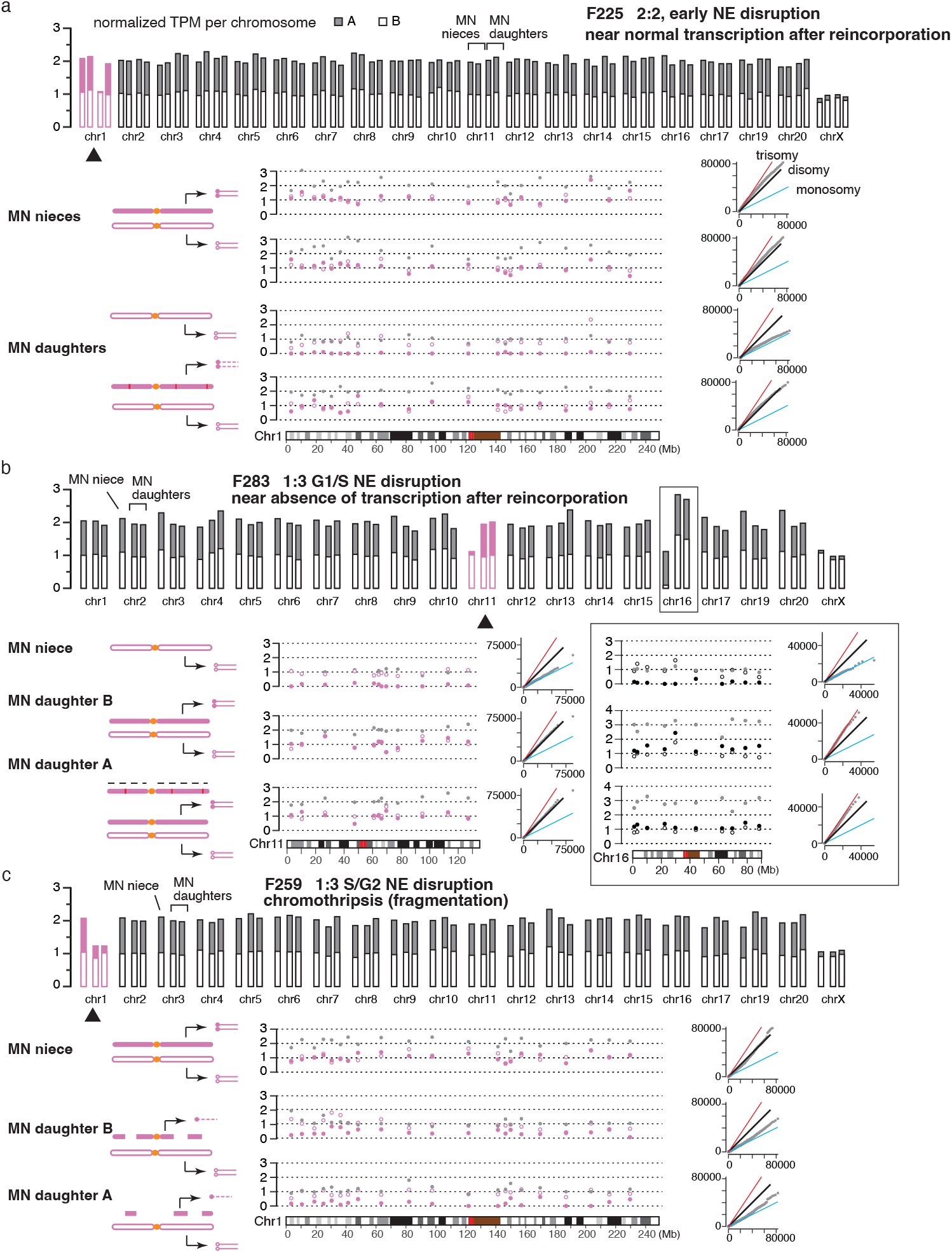
Examples of varying transcription of reincorporated MN chromosomes. (a) An example of near normal transcription of a reincorporated Chr.1 from a micronucleus in the F225 sample. The pattern of disomic transcription in both MN nieces and monosomic transcription in one MN daughter establishes the 2:2 segregation pattern. The presence of disomic transcription in the other MN daughter indicates close to normal transcription of the reincorporated Chr.1. (b) Left, an additional example of near complete absence of transcription of a reincorporated Chr.11 in the F283 sample. The pattern of monosomic transcription in one MN niece (the other was not captured) and disomic transcription in both MN daughters establishes the 1:3 segregation pattern. That both MN daughters display near disomic transcription indicates that the additional Chr.11 from the MN is not transcribed. Right, Chr. 16 is an example, where a chromosome was misegregated but, to the main nucleus. The monosomy evident in the niece means that one sister was monosomic and the other was trisomic. Both daughters of the trisomic sister show balanced expression of both haplotypes and near trisomic total transcription yield. Because both daughters share the trisomy, the missegregated copy of Chr. 16 cannot have been in the micronucleus. (c) An example of chromothripsis with partial incorporation of broken Chr.1 into both MN daughters in the F259 sample. The presence of disomic transcription in one MN niece (the other was not captured) and close to monosomic transcription in both MN daughters establishes the 2:2 segregation pattern. Although the average level of transcription in both MN daughters is near monosomic, we observe transcripts from the MN chromosome in both daughters in a partially reciprocal pattern (30-45 Mb and 200-230Mb); this pattern suggests reciprocal distribution of Chr.1 fragments into both daughters. Although chromosome fragmentation is expected to result in mirror-image pattern of DNA retention and loss, this pattern will not be consistently detectable at the RNA level for two reasons. First, the use of large bin sizes (50 genes or 10 Mb) necessitated by the sparsity of allelic expression signal may average out the oscillating pattern of DNA loss and retention. Second, the reincorporated DNA may undergo varying transcriptional recovery and the transcript abundance is not directly reflective of DNA copy number. Nonetheless, as chromothripsis is a frequent outcome of MN with S/G2 nuclear envelop rupture, we think chromothripsis is the most parsimonious explanation of the observed pattern. In total, we inferred chromothripsis to be present in five cases (out of 38 total), three showing deficient transcription recovery and two showing close to normal transcription when the transcription yield from both MN daughters are added together. As the inference of chromothripsis relies on the presence of MN haplotype-specific expression in both MN daughters, the inference of chromothripsis from single-cell RNA data inevitably underestimates the frequency of chromothripsis.

**Extended Data Figure 7.**
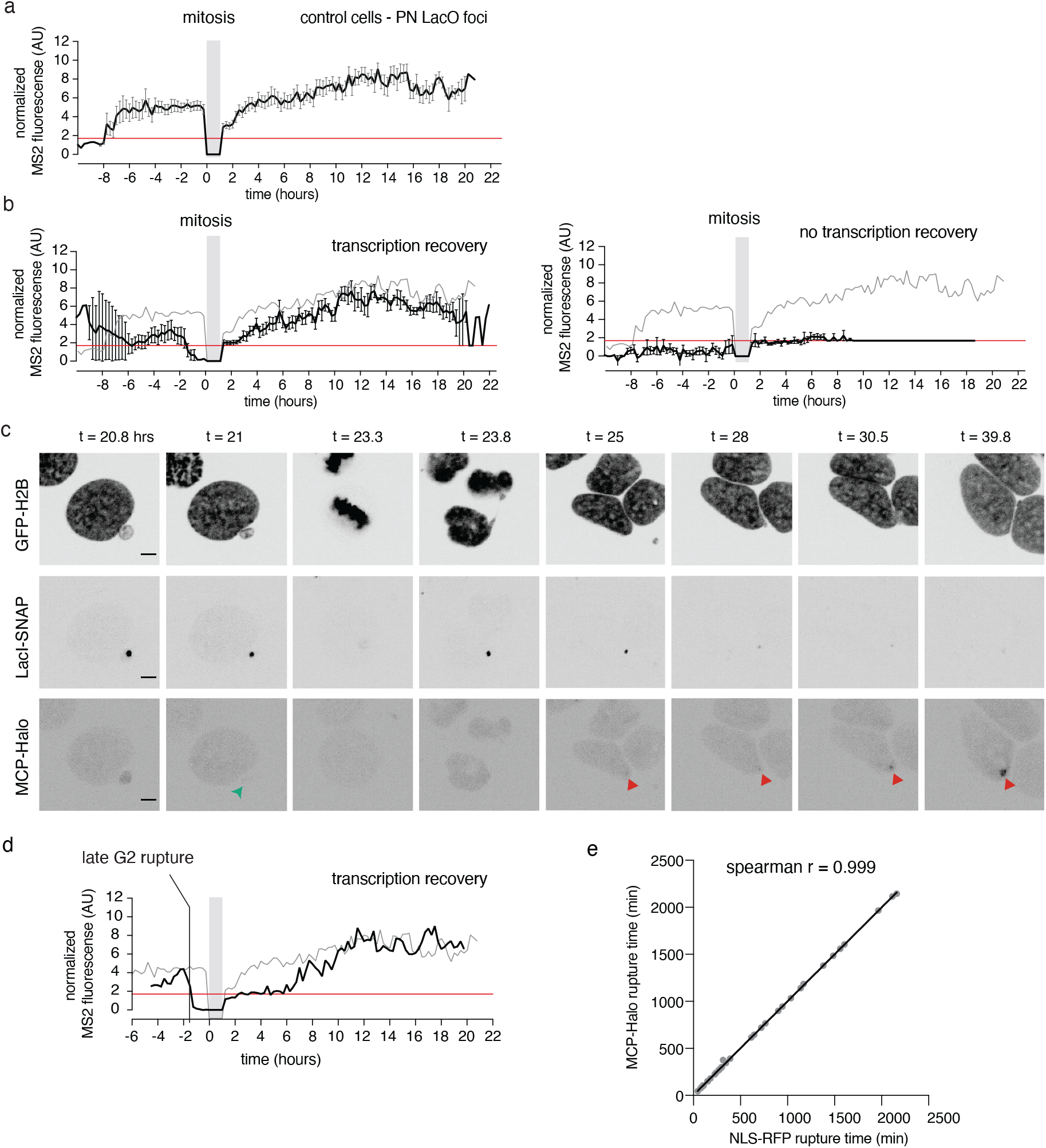
Analysis of nascent transcription from reincorporated micronuclei. (a) Control U2OS 2-6-3 reporters to assess nascent transcription of normally expressing reporters in the main nucleus. Fluorescence intensities of the MS2 signal (MCP-Halo) were measured from reporters that were in the main nucleus during both generation 1 and 2 (n = 23 LacI reporters). Grey bar: mitosis. Error bars: mean +/- SEM). Red line: minimum detectable intensity value of the controls (see Methods). (b) Aggregated data for samples using the U2OS 2-6-3 reporter to assess nascent transcription from reincorporated MN, similar to (a) and Fig. 2e,f. Fluorescence intensities of the MS2 signal (MCP-Halo) were measured from reporters that were in a MN in generation 1 and were then incorporated into a daughter nucleus in generation 2. Left, a subset of cells with MN that ruptured during generation 1 interphase and then recovered transcription after reincorporation into a daughter nucleus in generation 2 (n = 7 analyzed out of 19 similar cases). Note that prior to mitosis there is variable MS2 signal because of variable MCP-Halo accumulation in intact micronuclei and because of variability in the timing of MN NE rupture. Right, aggregate data for a similar subset of samples where the MN ruptured during generation 1 interphase and then displayed a generation 2 transcription defect after reincorporation (black line, n = 9 analyzed out of 20 similar cases). Grey line: mean intensity of the control reporters in main nucleus. Red line: minimum detectable intensity value of the controls. Note: for ease of visualization error bars (mean +/- SEM) are shown only for the experimental samples, not the controls. Also, when there was no detectable MS2 signal in the experimental samples, we assigned these samples to have the control minimal detectable intensity value (1.7, see Methods). This explains the complete overlap between the black and red lines after the 10 hour timepoint. (c) Images from a timelapse series for the experiment in (b), above. Green arrowhead: MN rupture. Red arrowheads: MS2 expression from the reporter after reincorporation into a daughter nucleus in generation 2. Time: hours post release from the G2 block. Scale bars 5 µm. (d) An example of a generation1 S/G2 MN rupture that recovered transcription after generation 2 reincorporation into a daughter nucleus. Performed and analyzed as in (a), above. (e) Validation of the method to assess MN rupture with the U2OS 2-6-3 reporter system. For all experiments with this reporter, loss of the general nuclear MCP-Halo signal (MCP-Halo contains an NLS) was used to determine the time of MN NE rupture. We verified that MCP-Halo signal loss from MN corresponds to RFP-NLS by two-color live-cell imaging in U2OS 2-6-3 cells expressing both MCP-Halo and RFP-NLS (n = 41 from four experiments; two-tailed Spearman’s correlation).

**Extended Data Figure 8.**
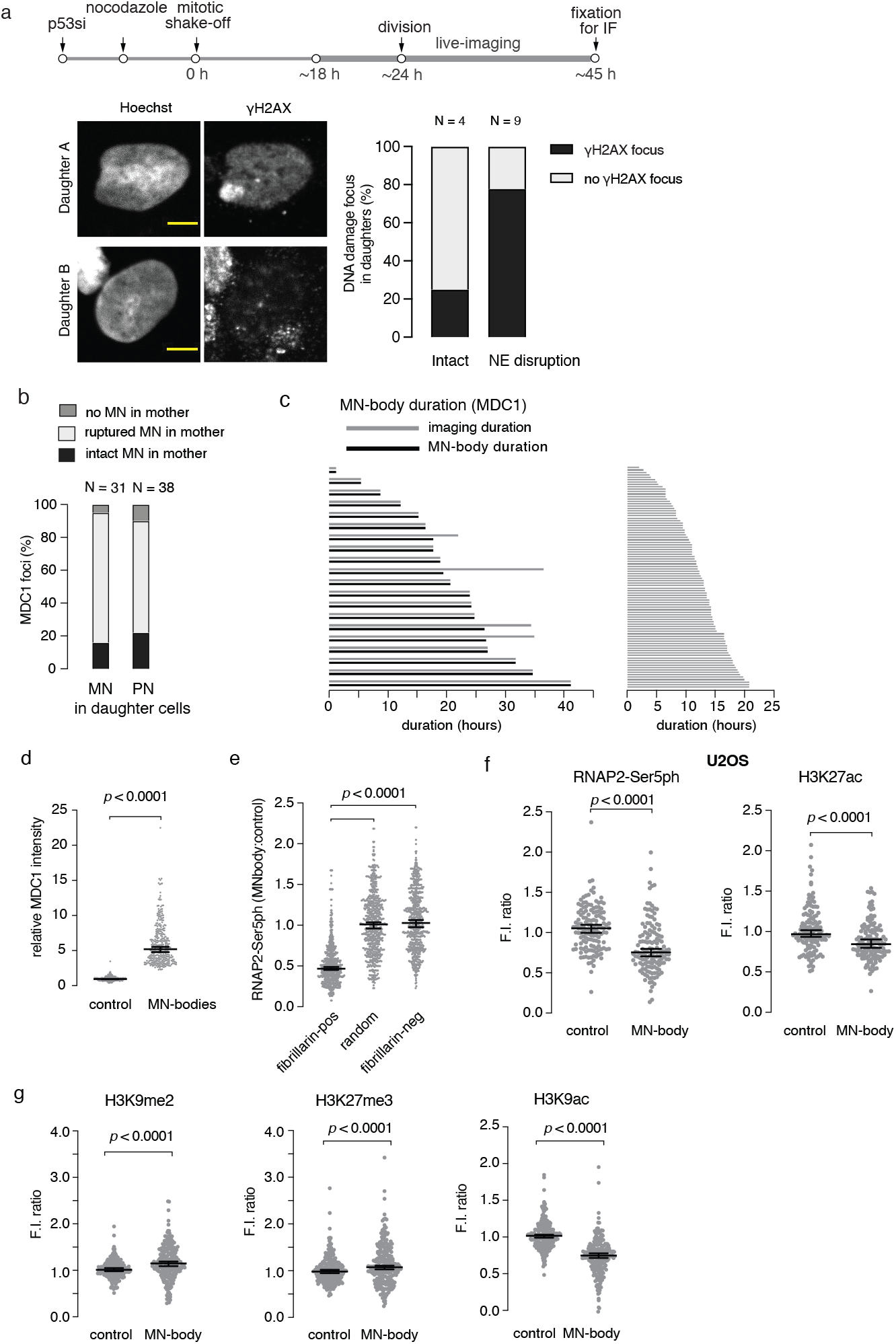
Characterization of MN-bodies. (a) Same-cell live-fixed experiment supporting the fixed imaging shown in Fig. 3b,c. Top, scheme of the experiment. MN were induced in RPE-1 cells, the MN fate was tracked with GFP-H2B and RFP-NLS (to visualize MN NE rupture, generation 1). After most cells progressed into generation 2, they were fixed and labeled to detect *γ*H2AX. Bottom left: representative images of a daughter cell pair, one with and one without an MN-body. Bottom right: summary of 13 cell pairs tracked and analyzed by the same-cell live-fixed experiments (from two experiments). Scale bars 5 µm. (b) Generation 2 MDC1-marked MN-bodies primarily derive from mother cells with MN. Live-cell imaging was performed as in Fig. 3a. and large cytoplasmic (persistent) or nuclear (reincorporated) MDC1 foci were identified. The movies were then analyzed to determine the frequency that cells with MN-bodies were derived from micronucleated mother cells and, if so, whether the MN ruptured or remained intact throughout generation 1. (c) Lifetime of MN-bodies. MN-bodies persist throughout most of the generation 2 interphase. Left: live cell imaging to assess SNAP-MDC1 marked MN-body lifetimes. Y-axis shows lifetime of individual MN-bodies, both the duration of imaging (light grey bars) and MN-body lifetimes (black bars). Note: that, with four exceptions, the MN-bodies persisted until the end of the imaging (see Fig. 3a). Right, lifetime of MN-bodies assessed by the same-cell live-fixed experiments from Fig. 3b,c, which has a larger sample size than the live-imaging on the left. Note: in this experiment, MN-bodies form at different times from the start of imaging and were followed until the time that imaging stopped, meaning that the duration of imaging and the duration of the MN-bodies are the same for each cell. (d) Distribution of signal intensities for MN-body by immunofluorescence staining for the endogenous MDC1. Performed and analyzed as in Fig. 3d (n = 341, from two experiments). Median with 95% CI. Two-tailed Mann–Whitney. (e) Determination of the background nuclear RNAP2-Ser5ph signal in nucleoli. We measured the background RNAP2-Ser5ph signal in nucleoli (fibrillarin positive), which should lack active RNA polymerase II, and in nucleus regions lacking nucleoli. These values were then normalized to the density of fluorescence intensity from a nuclear mask excluding the nucleoli-free. The fact that there is still measurable RNAP2-Ser5ph signal in the nucleoli means that we likely underestimate the extent of RNAP2-Ser5ph signal loss in MN-bodies (see Methods; n = 650, from two experiments; Kruskal-Wallis with Dunn’s multiple comparisons test). (f) Verification of low MN-body transcription and H3K27ac loss in U2OS cells. Performed and analyzed as in Fig. 3d (n = 138, from two experiments). (g) Reduced H3K9me2 (left) but not H3K27me3 (middle) or H3K9ac (right) in MN-bodies. Performed and analyzed as in Fig. 3d (n = 222, 234 and 244, left to right, from two experiments).

**Extended Data Figure 9.**
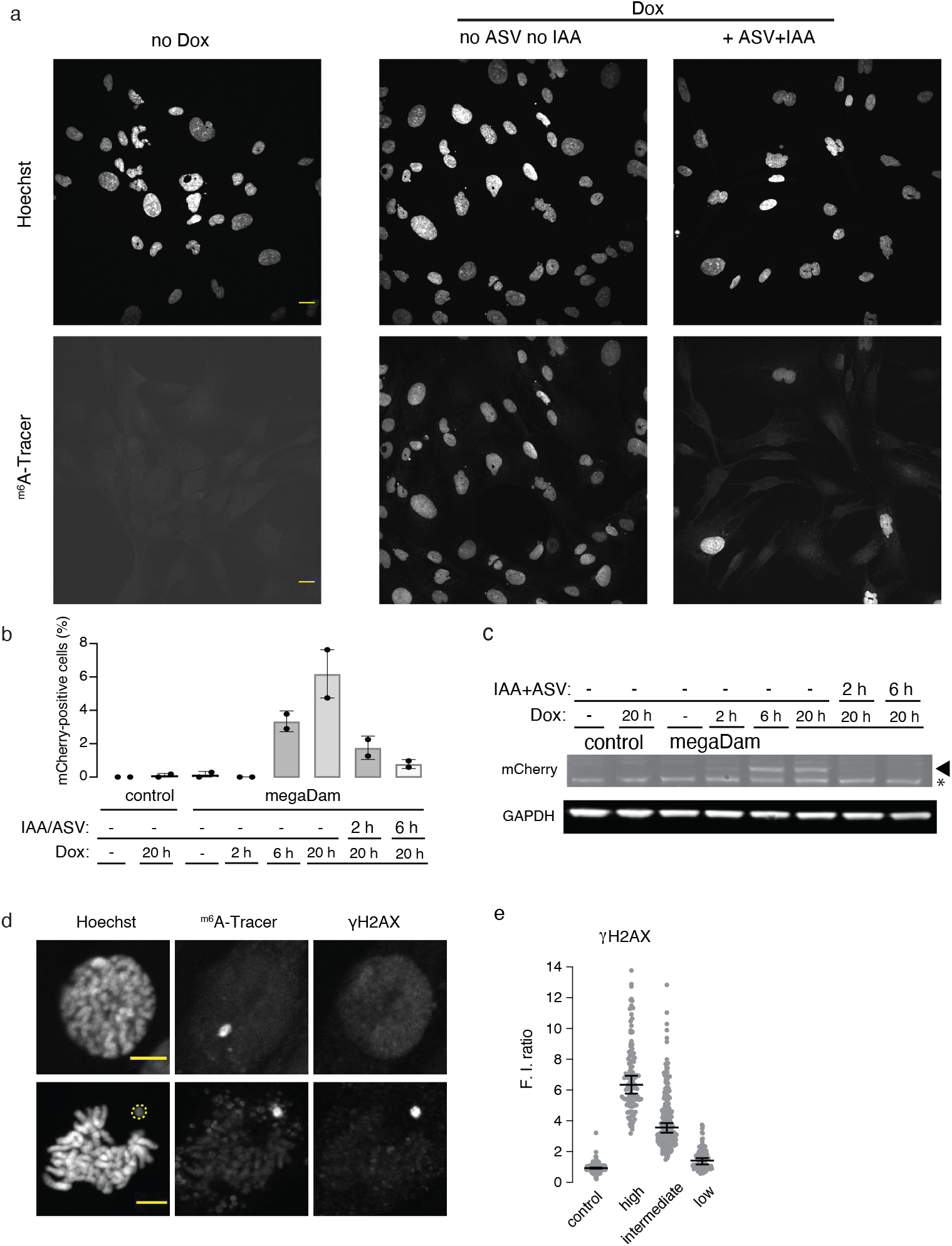
DamMN system characterization. (a) Validation of the DamMN system. Shown are representative single-focal plane confocal images of RPE-1 megaDam cells 45 h after post release from the CDK1-induced G2 block at the start of the experiment (see Fig. 4a and Methods). There is no m6A DNA methylation if megaDam transcript is not induced (left, no Dox); if megaDam is not degraded prior to mitotic entry, all primary nuclei show m6A DNA methylation because of labeling during mitosis (middle, Dox, no ASV no IAA), but primary nuclei are mostly not m6A methylated if megaDam is degraded prior to mitosis (right, Dox, +ASV +IAA). m6A methylation is visualized with the m6A-Tracer (see Fig. 4a and Methods; four experiments). Scale bars 20 µm. (b) Efficient induction and degradation of megaDam. FACS analysis to detect mCherry-tagged megaDam. All samples are unsynchronized RPE-1cells with or without megaDam, with or without megaDam transcriptional induction or megaDam degradation for the indicated periods of time. The controls are RPE-1 cells lacking the megaDam construct showing no background autofluorescence without or with Dox treatment. Shown is the percentage of cells expressing mCherry (PE channel, from two experiments). (c) Western blot to detect megaDam for the indicated samples corresponding to the experiment shown in (b), above. Shown is a cropped image of a gel from the region at the megaDam molecular weight (130 kDa). Note: the *α*-mCherry Ab detects non-specific background bands, but megaDam is readily distinguished from these background bands (two experiments). * indicates a background band. (d) Specific labeling of MN chromosomes in mitotic cells (two examples) using the DamMN system. Top shows an MN chromosome in a prometaphase cell lacking DNA damage. Bottom shows an MN chromosome in a metaphase cell with DNA damage. Note that the MN chromosome is less condensed during mitosis, as has been previously described ^48^. Performed as described in Fig. 4a. Yellow dashed line: an MN chromosome positive for *γ*H2AX and m6Tracer (n = 4 experiments). Scale bars 5 µm. (e) Control for Fig. 4c showing the distribution of MN-body *γ*H2AX FI units relative to the general nuclear background (lacking nucleoli). The MN-body region of interest corresponds to the m6A-Tracer signal (see Methods). The *γ*H2AX low MN-bodies were designated if the total area of *γ*H2AX positive pixels (>3SD above background, see Methods) occupy less than 21% of the MN-body area (corresponding to the bottom quartile of *γ*H2AX positive MN-bodies). The designation of *γ*H2AX intermediate MN-bodies was between 21% and 65.7% of the MN-body area (the middle two quartiles), and *γ*H2AX high was *>* 65.7% of the MN-body area (the top quartile of MN-bodies, four experiments). Error bars: median with 95% CI.

